# Getting Blood From a Stone: Improving Brain-Behavior Inferences Without Brain Data

**DOI:** 10.1101/2021.01.21.427334

**Authors:** David Halpern, Todd Gureckis

## Abstract

In recent years, the cognitive neuroscience literature has come under criticism for containing many low-powered studies, limiting the ability to make reliable statistical inferences. Typically, the suggestion for increasing power is to collect more data with neural signals. However, many studies in cognitive neuroscience use parameters estimated from behavioral data in order to make inferences about neural signals (such as fMRI BOLD signal). In this paper, we explore how cognitive neuroscientists can learn more about their neuroimaging signal by collecting data on *behavior alone* and using alternative estimators designed to leverage this information. We demonstrate through simulation and mathematical derivations that knowing more about the marginal distribution of behavior can improve inferences about the mapping between cognitive processes and neural data. We analyze the magnitude of this benefit, finding that it depends on the desired estimand and several underlying study parameters. While in many cases the absolute gains in precision can be modest, our results demonstrate that, in realistic settings, additional behavioral data can lead to the same improvement in the precision of inferences more cheaply and easily than collecting additional data from subjects in a neuroimaging study. This means that when conducting a neuroimaging study, researchers now have another knob to turn in a design analysis: the number of subjects collected in the scanner and the number of behavioral subjects collected outside the scanner (in the lab or online).

## 1 Introduction

One factor which limits progress in human cognitive neuroscience is statistical precision and power (Cremers et al., 2017; Munafò et al., 2019; Yarkoni, 2009). Many reported effects are small relative to the noise in neuroimaging signals, a problem compounded by the small samples sizes used in most studies. Within a hypothesis testing framework, a lack of precise estimates of some measurements can make it difficult to replicate reported effects. For example, even quite strong, known effects such as the relationship between motor movements and fMRI BOLD activation in sensorimotor regions can be unlikely to reach significance in current standard sample sizes of around 25-30 subjects (Poldrack et al., 2017). Having lower precision also means that any estimated effect that is significant will also have a high probability of the estimate being much greater in magnitude or even having the wrong sign compared to the true effect (Cremers et al., 2017; Gelman & Tuerlinckx, 2000; Yarkoni, 2009). However, due to a combination of factors (including lack of sufficient funding for large sample studies), research in human cognitive neuroscience tends to be under-powered in many cases (Poldrack et al., 2017).

Several remedies have been proposed for this situation, the most obvious being to simply collect more neuroimaging data (Munafò et al., 2019; Poldrack et al., 2017). Because the funding available to most labs is relatively small (compared to the cost of a large scale neuroimaging study), this might require a move towards working in larger consortia around critical topics. Another approach might be to leverage the power of publicly available open datasets, which can allow for meta-analyses across several smaller experiments (Poldrack et al., 2017). However, increasing sample sizes is not the only option for dealing with a lack of precision. For example, increasing the signal-to-noise ratio of neuroimaging measures by creating more detailed statistical models of the neuroimaging signal (Lindquist et al., 2009), improving experimental design (Durnez et al., 2017) or even improving the measurement process itself (Feinberg & Yacoub, 2012; Lombardo et al., 2016) are all measures that are likely to help. In addition, it has been suggested that a focus on computational cognitive modeling and behavior can also allow for extracting more signal (Krakauer et al., 2017; Palmeri et al., 2017) by creating better models of the underlying cognitive and neural processes. In that theme, this paper explores a potentially under-studied option for improving the statistical precision of human cognitive neuroscience studies: collecting additional behavioral data without neural recordings.

We begin by noting that many common statistical analyses in cognitive neuroscience involve behavioral data, typically collected from the subjects who also provided neural data. In one common scenario, the interest is in the relationship between individual differences in behavioral traits and more static measures of neural activity, such as performance in a task and functional connectivity (e.g. Rosenberg et al., 2016). In another typical analysis, researchers are interested in how within-subject changes in a cognitive state variable over time relate to changes in the neural signal from a particular brain region, such as the prediction error in a reinforcement learning task (e.g. O’Doherty et al., 2003). One important, but often overlooked, point is that these behavioral measurements (e.g.. survey questionnaires, behavioral responses to stimuli, summaries of task performance or estimates of cognitive model parameters) are themselves noisy estimates of an individual’s underlying cognitive processes and the distributions of these patterns that exist in the population. Thus our ability to infer relationships between brain and behavior is limited by *sampling error* and *estimation error*. Sampling error is a result of the fact that a small sample may not be representative of the population, so estimates of the relationship based on the sample may be far from the true relationship. A small sample therefore prevents us from being confident about the value of a parameter in a statistical model. Estimation error is a form of measurement error that arises because researchers cannot measure latent cognitive variables directly and instead use models to estimate their values. The relationship between neural recordings and cognitive variables measured with error will necessarily not be as strong as the relationship involving the true values, a notion sometimes known as regression dilution or attenuation. Ignoring these two sources of error can prevent us from learning as much as possible from neural data.

In the following, we demonstrate that one way to reduce both sources of error and improve statistical estimates of the relationship between the brain and behavior is to collect behavioral data from subjects *without collecting corresponding neuroimaging data*. This idea is somewhat counterintuitive in light of the fact that most common approaches in neuroimaging analyses assume that the only relevant data is from individuals whose neural activity was recorded. However, behavioral data (especially data collected over the Internet) is often significantly less resource-intensive to collect than neuroimaging data, in terms of money and the time and effort of both researchers and human subjects. We show how using modern statistical methods that can leverage information from the additional behavioral data can help mitigate the two types of error. To anticipate one consequence of our analysis, we are able to provide a cost-benefit analysis (in terms of statistical power and precision) of collecting data from subjects with only behavior against collecting more expensive additional data from subjects with both behavior and neural recordings. In some cases, due to its relatively low cost, collecting more behavioral data instead of additional neuroimaging data can be a more effective use of resources. This result has implications for designing studies in cognitive neuroscience both prospectively and retrospectively (i.e., the re-analysis of archived open-science data sets.)

### 1.1 Notation

Throughout the article, we will discuss a situation where we have collected both neuroimaging data (**y**) and behavioral data (**x**) from *N*_*xy*_ participants and we will ask what the benefit is of collecting an additional *N*_*x*_ participants who only contribute behavioral data (from the same measurement as in the original experiment). We will always refer to constants, like the number of participants, with capital letters (*N*). In addition, we refer to data vectors with lower case letters, **x** and **y**. Estimated latent data variables from cognitive models (defined below) will be referred to as *c* when they vary within-person (state variables) and *θ* when they only vary across people (trait variables).

### 1.2 Common paradigms for relating brain and behavior in neuroimaging studies

For the purposes of articulating our thesis, it will be useful to describe two common types of analyses used in cognitive neuroscience that use behavioral data to interpret neural signals and lay out some notation for the rest of the paper. While this is by no means an exhaustive list, we claim that many analyses can be understood as falling into two broad categories.

#### Regression (Conditional Distribution)

In many cognitive neuroscience studies, researchers are interested in how the conditional distribution of neural measures changes as a function of variation in cognitive states induced by a task. For instance, researchers might be interested in regions of the brain that track whether or not it is viewing a face (Kanwisher et al., 1997), the valence of the currently presented stimulus (Chikazoe et al., 2014) or the prediction error in a reinforcement learning model of decision making (O’Doherty et al., 2003). More formally, a subject *i* presented with a set of stimuli **s**_**i**_ that occur on trials (or times) *t* ∈ *{*1…*T*_*i*_*}*. Each stimulus presentation *s*_*it*_ is assumed to induce some latent cognitive state *c*_*it*_ that is associated with some neural activity *y*_*it*_. Researchers can identify neural signals related to the cognitive states by fitting a regression model where

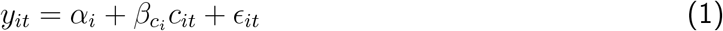

where *ϵ*_*it*_ is uncorrelated noise (for the purposes of this discussion, we ignore features of particular signals like the fMRI BOLD hemodynamic response). However, this is not quite so straightforward because researchers need to derive estimates of the cognitive state for these analyses. The most common assumption is that the relevant states are closely tied with objective features of the stimulus, such as the color of an image or experimental conditions like the reward offered for a choice, so *c*_*it*_ can reasonably be replaced by the stimulus feature or experimental condition itself.

In other cases, the cognitive states of interest are tied to subjective features of the stimulus, such as the valence associated with *s*_*it*_ or the typicality of *s*_*it*_ for a particular category. In some cases, researchers collect behavioral ratings *b*_*it*_ which depend on these latent states. Researchers then assume that the ratings come from a cognitive process *C* such that *b*_*it*_ ~ *C*(*c*_*it*_, *s*_*it*_) and obtain the *c*_*it*_ from that model. In the simplest case, this can mean simply using the valence ratings given by subjects (Chikazoe et al., 2014) or using the average typicality from a group of subjects tested separately (Wilson-Mendenhall et al., 2015).

More recently, “model-based” neuroimaging analyses have investigated cognitive states that are involved in a particular cognitive process *C* such that in addition to the current stimulus *s*_*it*_, *c*_*it*_ may also depend on *θ*_*i*_ are latent cognitive “trait” variables that do not vary over the relevant time as well as the past states 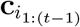. Researchers can then use the behavioral data to fit the model *b*_*it*_ ~ *C*(*θ*_*i*_, *c*_*it*_, *s*_*it*_) and obtain the relevant *c*_*it*_. These states may be, for instance, the reward prediction error or predicted value in a reinforcement learning model of sequential decision making (O’Doherty et al., 2003).

Because the latter two types of analyses involve behavioral data, those will be our focus in this paper. In particular, we will focus on the model-based case but nearly all of our analyses will also apply to the second case.

Modern multivariate pattern analysis techniques (Haxby, 2001; Polyn, 2005) invert this regression and try to predict the cognitive state from neural signals. Typically, this means predicting stimulus properties or experimental conditions but in some cases the cognitive state will be inferred from behavioral data (e.g. Kragel & Polyn, 2016; Mack et al., 2013). While the example analyses in the rest of the paper do not directly apply to this setting, similar principles of collecting additional behavioral data may hold there as well.

#### Correlation (Joint Distribution)

In other cases, researchers are interested in the joint distribution of behavioral measures and neural measures (often across subjects). For instance, is there a relationship between individual differences in behavior and individual differences in static measures of the brain? Researchers often use a direct measure of a subject’s behavior *x*_*i*_ such as answers to a questionnaire (Treadway et al., 2013) or performance on a behavioral task (such as a sustained attention task Rosenberg et al., 2016). More recently (e.g. Homan et al., 2019), researchers have also used cognitive “trait” variables which can be related to behavioral data through cognitive models (i.e. *θ*_*i*_ above). Static measures can include measures like functional connectivity or average resting state activity that do not depend on time. Typically we only get one *y*_*i*_ per subject and the measure is not always collected simultaneously to the task where *x*_*i*_ is recorded. In this setting, researchers are typically interested in characterizing the joint distribution of neural measures *y*_*i*_ and behavioral (*x*_*i*_) or latent cognitive (*θ*_*i*_) trait variables. In particular, if the neural measures and behavioral variables can be approximated by a bivariate normal, researchers want to find neural measures where the correlation *ρ* is non-zero.

Another common analysis that involves correlation is Representational Similarity Analysis (Kriegeskorte, 2008). In these analyses, researchers correlate the similarities of the neural response to pairs of stimuli with the similarities of the two stimuli according to a model. Most often, these models are created based on stimulus properties but they are sometimes derived from behavioral ratings (Bruffaerts et al., 2013; Chikazoe et al., 2014). In these cases, our analyses about the benefits of additional behavioral data will apply as well.

We argue that these two categories of research design and inferential techniques cover a vast array of studies in cognitive neuroscience and neuroimaging in particular ^1^. Each of these analyses can be thought of as attempting to learn about the *conditional* distribution of neural signals given behavioral data (**regression**) or the *joint* distribution of neural and behavioral data (**correlation**). The focus of this paper ask whether we can improve inferences about the relationship between brain and behavior in these two types of analyses by learning more about the *marginal* distribution of behavioral data, that is, collecting more behavioral data without neural data.

## 2 Two Sources of Error in Neuroimaging Analyses

As the Introduction laid out there are two key sources of error in neuroimaging analyses. In this section, we describe each of the two sources of error in more detail as well as how they can affect the precision of inferences about the relationship between neural signals and behavior.

### 2.1 Sampling Error

In a **correlation** analysis, we want to know what *ρ* is in the general population but estimate it by using the sample correlation *r*. With a small number of subjects, *r* can be quite far from the true *ρ*. An approximate confidence interval for *ρ* can be constructed as arctanh 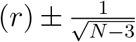, where *N* is the number of subjects or, more generally, the number of data points used to estimate *r* (Fisher, 1915). As this makes clear, the precision of the estimate (and thus finding a *ρ* that we can say is non-zero with confidence) is constrained by the size of our sample. In many neuroimaging studies this is often relatively small because neuroimaging data is expensive and time consuming to collect. In Figure 1, we show an example of where the smaller sample size can lead to noisy estimates and larger error bars. ^2^ Sampling error can also be an issue in **regression** analyses as well. However, because in that case we are only interested in the conditional distribution of the neural signals, we cannot reduce sampling error by collecting more behavioral data so do not discuss it here.

**Figure 1:**
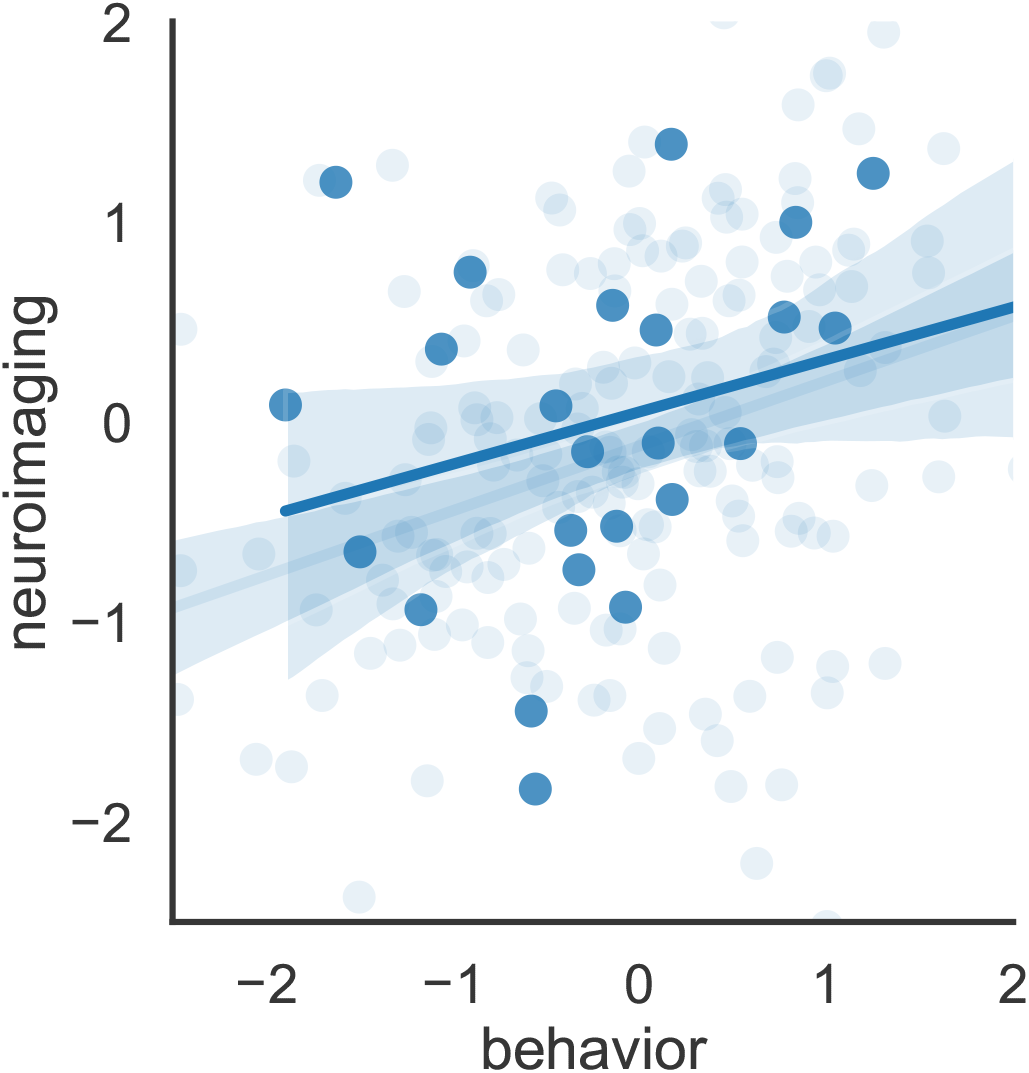
25 samples (dark blue) from a multivariate Gaussian distribution are overlaid over an additional 100 samples (light blue) from the same distribution. Regression lines are plotted showing how estimates from smaller samples can add significant variance to the estimated correlation. Error bars show that we cannot rule out a zero correlation in the smaller sample.

**Figure 2:**
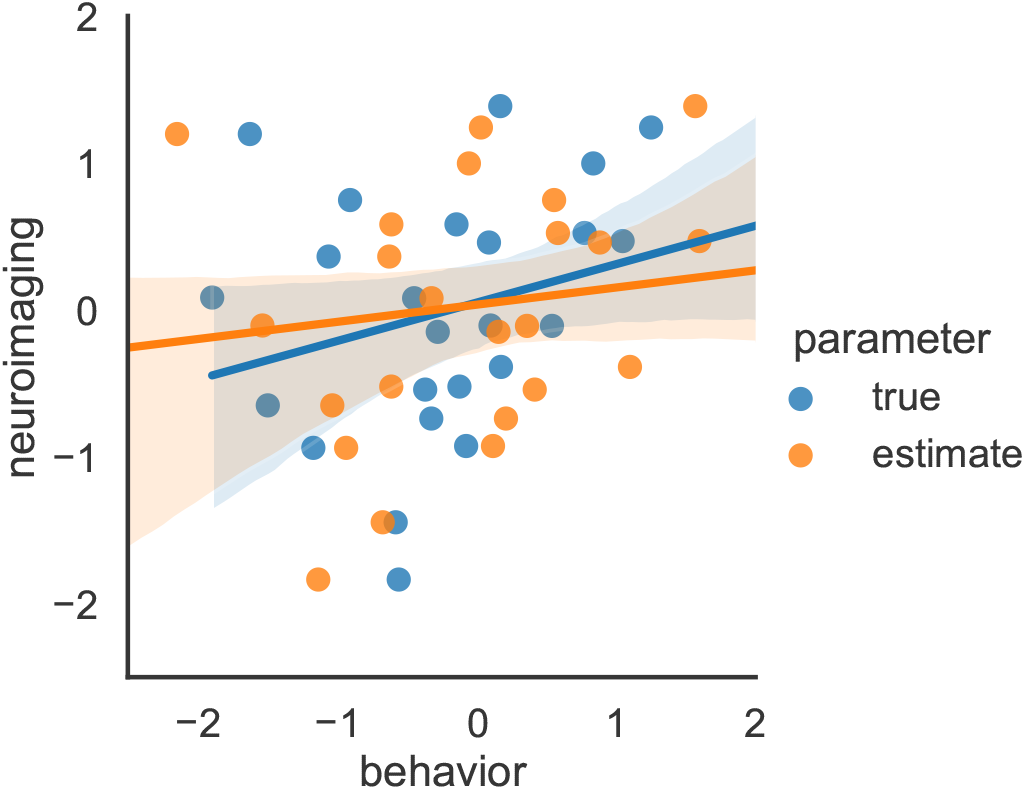
Here we show a case where the latent behavioral measure is estimated with error. The correlation with the estimates is lower than it would be with the true latent behavioral measure.

### 2.2 Estimation Error

In the **model-based regression** analysis described above, we want to find a set of neural signals that have a linear relationship with a subject’s latent cognitive states **c**_*i*_. ^3^ The true model we are interested in is

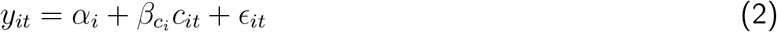

where in particular we are interested in which neural signals have a significant 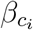 parameter (typically, at the group level). In practice, because we don’t have direct access to *c*_*it*_ we typically replace it with estimates *ĉ*_*it*_ by fitting the model *C*(*θ*, **c**_*it*_, **s**_*it*_) to the behavioral data *b*_*it*_. It is typically assumed that the model doesn’t perfectly account for behavior and that there is some variability in responses beyond what can be predicted by the model. For instance, when fitting a reinforcement learning model, a decision noise parameter is often used to deal with variability beyond what can be accounted for by the latent representations in the model. Because we don’t have infinite data, this typically means that several sets of parameters could account for the data even if the model is well identified. In other words, the parameters **ĉ**_*it*_ are estimated with some uncertainty. This may seem obvious to many readers but standard techniques that are typically used in cognitive neuroscience do not easily allow this uncertainty to be taken into account in the above regression (c.f., Ly et al., 2017; B. M. Turner, 2015; B. M. Turner et al., 2013, 2017). Instead of the model above, researchers often simply plug in the estimated **ĉ**_*i*_ into the regression, i.e.

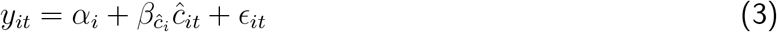

In this model, the value of the slope estimate using the estimated latent variables, 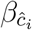, will not necessarily be the same as when using the true parameter 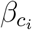. In fact, it can be shown that on average, the least squares estimate of 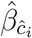 will be

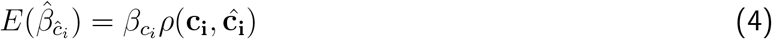

where *ρ*(**c**_**i**_, **ĉ**_**i**_) is the correlation between the true **c**_**i**_ and the estimates **ĉ**_**i**_ (Wilson & Niv, 2015). This is perhaps not always relevant since the neuroimaging signal can often only be known up to a multiplicative constant. However, Wilson and Niv (2015) also showed that, as long as the estimates are uncorrelated with the regression error term *ϵ*_*it*_, the expected *t*-statistic for the 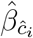 is

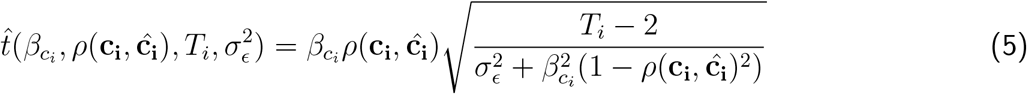

where *T*_*i*_ is the number of trials from subject *i* used to estimate the regression and 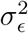 is the variance of the residuals *ϵ*_*it*_, due to noise in the neuroimaging signal. We can see that this *t*-statistic gets closer to 0 as *ρ*(**c**_**i**_, **ĉ**_**i**_) gets further from 1. A smaller t-statistic means that the standard error (or any multiple of it such as a confidence interval) is more likely to overlap with 0. Thus having a lower correlation between the estimates and the true values, or, in other words, greater estimation error, can make it harder to find relevant signals in the typical regression setting.

Estimation error can also be relevant for **correlation** analyses looking at correlations between latent cognitive trait variables and neural signals. If we estimate the trait variable *θ*_*i*_ from behavioral data, Katahira (2016) showed that the correlation between estimates 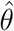 and structural neural signals **y**, can be decomposed as

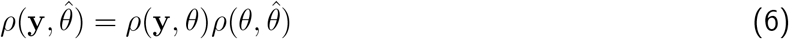

Therefore, the correlation is upper bounded by the true correlation between the neural signals and latent trait variables *ρ*(**y**, *θ*) and will decrease as the correlation between the estimates and the true values 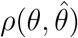 decreases (or as estimation error increases). Because the size of confidence intervals for a correlation coefficient only depend on the total number of subjects, the intervals for a correlation with a noisier estimate will be more likely to overlap with zero. This means that again, greater estimation error will make it harder to find relevant signals.

## 3 The Benefits of Additional Behavioral Data

Having laid out why these two errors can deteriorate our inferences, we now demonstrate how collecting additional behavioral data can help. In addition, for each case, we lay out a tradeoff between collecting cheaper behavioral data and more expensive imaging data in terms of how much each improves our inferences. These analyses show in which cases it may actually be more valuable to collect behavioral data than neuroimaging data, even if the main object of interest is the relationship between brain and behavior. To preface, section 3.1 demonstrates how little known estimators are able to make use of additional behavioral data to reduce sampling error for correlation analyses. We then follow with simulations to understand where this estimator improves over standard ones. Section 3.2 then documents how the use of hierarchical modeling can allow for combining datasets and improving estimation for common regression analyses as well as correlation analyses. Further analytical work shows the benefits of decreasing estimation error for making group-level inferences about the relationship between cognitive processes and neural signals.

Critically, these are two *distinct* methods by which behavioral data can decrease sampling error and estimation error. For each, we show why each method works and then, using a combination of analytic derivations and simulations, investigate the tradeoff between collecting additional behavioral data vs. neuroimaging data when using these methods. As mentioned above, regression analyses commonly used in cognitive neuroimaging are not affected by the population variance of the behavioral variable. However, in the case of correlational analyses, or analyses of the full joint distribution of neural and behavioral measures more generally, both methods could be used, potentially in combination. The following table summarizes these situations more succinctly.

**Table 1:**
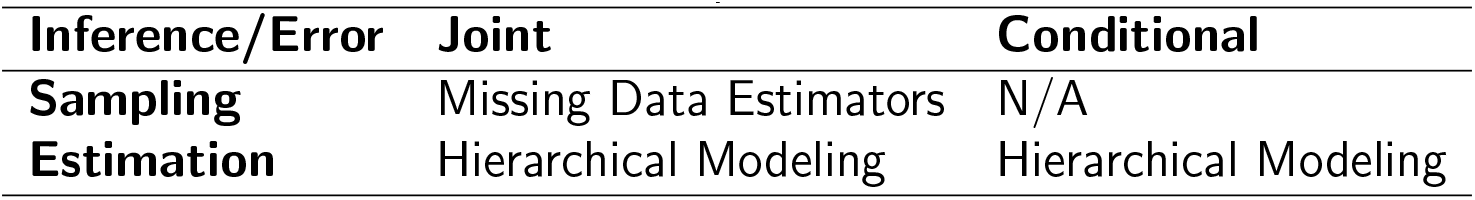
Proposed Methods.

### 3.1 How to Decrease Sampling Error: Behavioral Data as Neuroimaging Data with a Missing Variable

Typically we are interested in generalizing beyond our sample and characterizing the joint distribution of neural signals and behavioral variables in the population. However, as mentioned, most neuroimaging studies are limited by getting a relatively small sample. One somewhat counter-intuitive notion from the field of statistics focused on missing data is that we can often get better estimates of a parameter in the population (in terms of a lower mean squared error) by including data points that do not have every variable recorded in our analysis (Little & Rubin, 2019). This is because we are often assuming that variables have some joint distribution. If the correlation between two variables is not zero then we can know something about the value of what a missing variable must be from knowing the value of the other variable. If we have collected neural and behavioral data from *N*_*xy*_ subjects and only behavioral data from an additional *N*_*x*_ subjects, Anderson (1957) showed that the maximum likelihood estimates of the mean and standard deviation of **y**, as well as the correlation between **y** and **x**, *ρ*, are not the same when we include all of the data points as they would be if we only include the data with all variables recorded (the complete data). We can see this by rewriting the likelihood for a bivariate normal. If

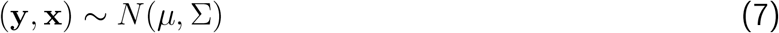

where *µ* = (*µ*_**y**_, *µ*_**x**_) and 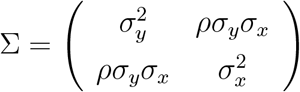 then, we can factor the likelihood as

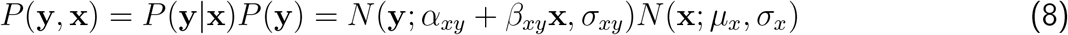

The advantage of writing the joint distribution as we did above is that the first term is just the regression of **y** on **x** which is only defined for the data points that have both variables recorded. Therefore, we can get the maximum likelihood likelihood estimate for *α*_*xy*_, *β*_*xy*_ and *σ*_*xy*_ using standard least squares regression estimates with the *N*_*xy*_ data points. Maximizing the likelihood of the second term only depends on the **x** data so we can use standard estimates of the mean and variance using all of the data. We can write the correlation as

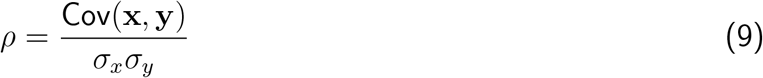

Then plugging in the maximum likelihood estimates 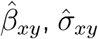 and 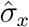, we get Anderson’s factored likelihood estimator for *ρ* using all of the data:

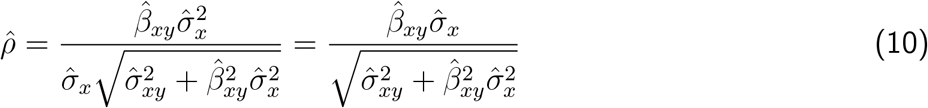

Because the maximum likelihood estimate of *σ*_*x*_ using all *N*_*xy*_ + *N*_*x*_ data points will not in general be the same as only using the *N*_*xy*_ data points, the maximum likelihood estimate of *ρ* using all the data will not be the same as using only the complete data.

In the Appendix, we show that there is a simple relationship between the standard Pearson *r* and Anderson (1957)’s 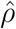. Specifically, if 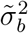 is the estimated standard deviation using only the *N*_*xy*_ subjects with neuroimaging data and 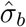 is the estimated standard deviation using all *N*_*xy*_ + *N*_*x*_ subjects then

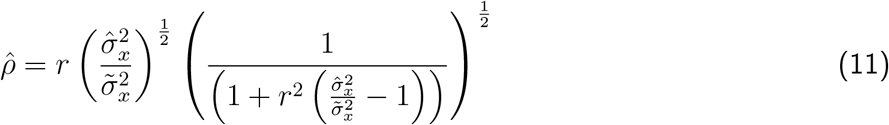

To our knowledge, we are the first to show that 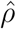, only depends on the correlation estimate in the complete cases, *r* and the ratio of the variance of the behavioral variable in the complete cases and the full data set, 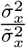. This form makes clear how the new information on the marginal distribution of the behavioral data contributes to the estimated measure of association. In figure 3, we plot how 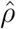 changes as a function of these two variables, with the variance change plotted on the log scale. In general, if the variance increases when including the additional data, the correlation estimate increases and visa versa.

**Figure 3:**
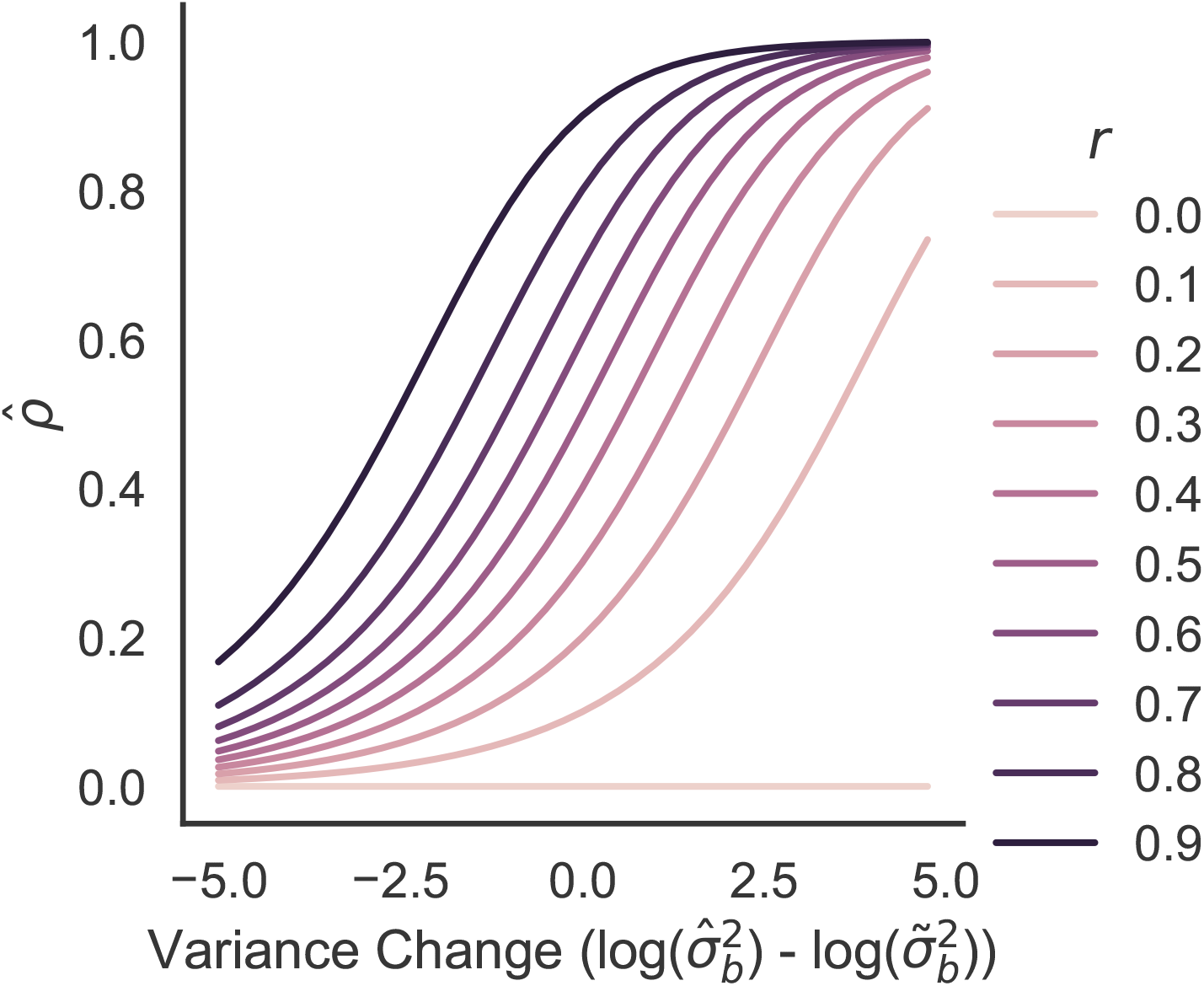
Anderson (1957) estimate 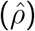 of the correlation using all of the data as a function of Pearson’s *r* using only *N*_*xy*_ complete cases and the difference in the log of the variance of the behavioral data in the *N*_*xy*_ complete cases and all *N*_*x*_ cases. In general 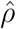 is modified up or down by the relationship between the variance of the behavioral data on the full data set and that for the complete cases.

#### 3.1.1 When does it work?

Garren (1998) showed analytically that asymptotically, the maximum likelihood estimate 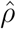 using all the data will have lower error than estimates for *r* using only the complete data. This result suggests that there can be cases where, if one variable is more expensive to collect than the other one (for instance neuroimaging data and behavioral data), the same quality of estimates of the correlation can can be achieved for lower cost by collecting some data with the more expensive variable missing (Hocking & Smith, 1972). However, the result is an asymptotic result, meaning that it is only guaranteed to apply in the limit of infinitely large data sets. It is therefore still unclear when collecting more behavioral data will be better than collecting more neuroimaging data in practice. In the following, we conduct a simple simulation to explore this tradeoff.

Following the description above, suppose we have collected a behavioral variable (**x**) and a neuroimaging variable (**y**) from *N*_*xy*_ subjects. In addition, we collected just behavioral data from *N*_*x*_ subjects. For simplicity, we assume that we have a single behavioral measure and a single neuroimaging measure from each subject.

In order to specify the generative model with missing data, we can create a matrix of data **XY** from our neuroimaging subjects, an *N*_*xy*_ *×* 2 matrix with **y** as the first column and 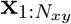 as the second. The data matrix is generated for *i* ∈ [1, *N*_*xy*_] according to

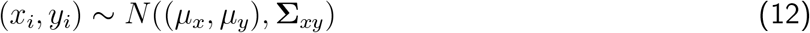

where 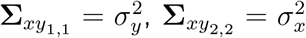 and 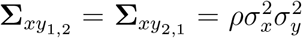. The additional behavioral subjects then have distribution

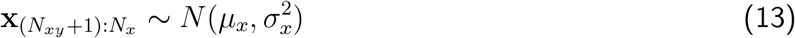

For the purposes of our simulation, we set *µ*_*x*_ = *µ*_*y*_ = 0 and 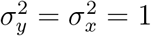.

Having generated data according to the above model, we now compare two ways of estimating the correlation, standard Pearson’s *r* using only data with both variables and the Anderson (1957) estimate of the correlation using all of the data.

As an example, we first compare estimates at *ρ* = .5 for a study with 25 neuroimaging subjects. We then compare Pearson’s *r* and Anderson’s 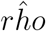 for five values of the number of additional behavioral subjects (0, 25, 50, 100, 250, 500 and 1000). Finally, to compare the benefit of the additional behavioral data to collecting an additional neuroimaging subject, we compare all estimates to Pearson’s *r* with one additional neuroimaging subject. We simulate 1,000,000 datasets and estimate the root mean squared error of all of these estimates averaged across all simulations. We can see in Fig. 4 that the Anderson estimator quickly improves over obtaining an additional neuroimaging subject.

**Figure 4:**
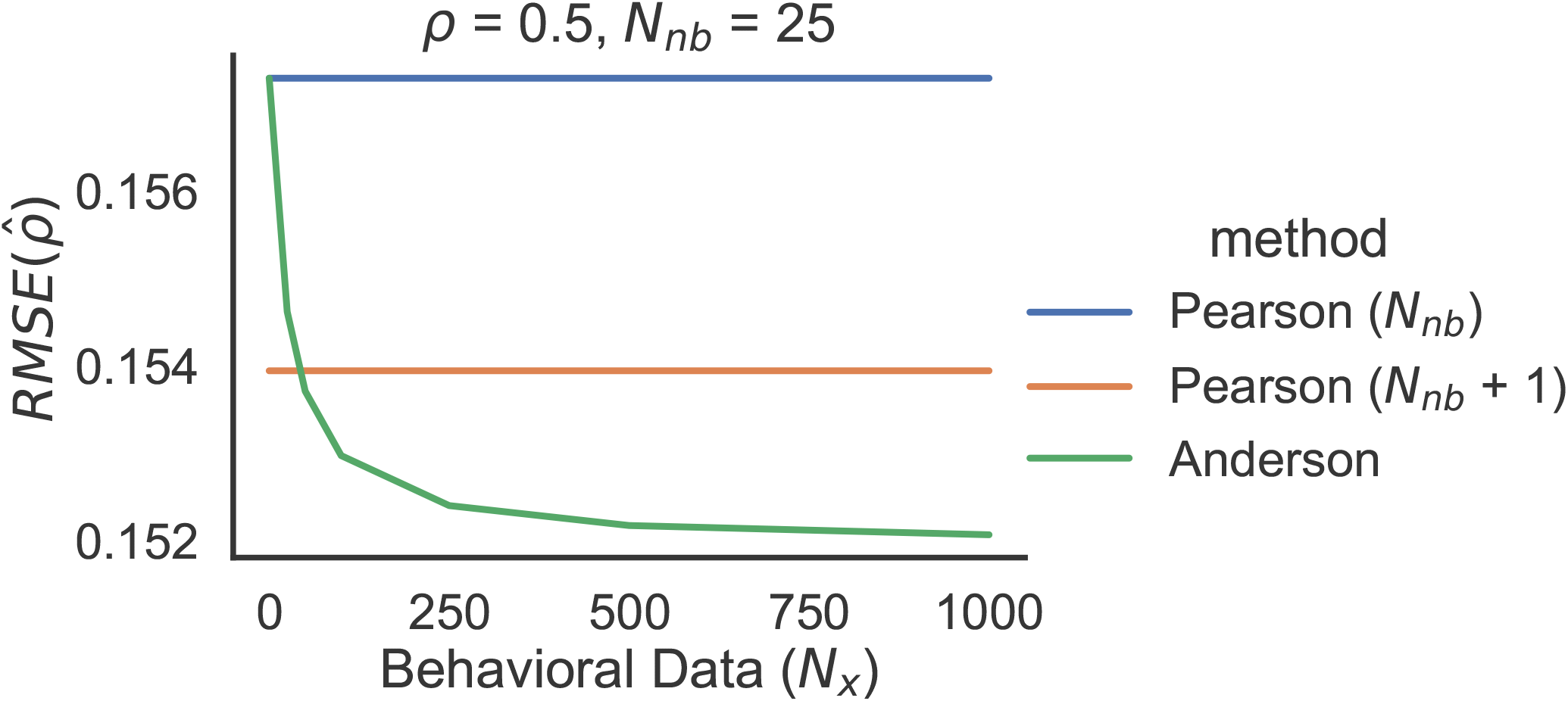
Root mean squared estimation error, averaged over 1,000,000 simulated datasets, for the estimate of *ρ* from the two estimators as a function of the true correlation and number of additional behavioral subjects *N*_*x*_ for *ρ* = .5 and *N*_*xy*_ = 25. We also compare to estimates from a neuroimaging dataset with *N*_*xy*_ + 1 subjects. The orange line is below the blue line showing the improved estimate that comes from one additional neuroimaging subject. The curved green line shows the Anderson (1957) 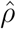 changes in the estimate as the number of addition behavior only subject are added to the analysis.

Of note here is that collecting neuroimaging data can be quite expensive. For instance in 2022, collecting fMRI data at New York University’s Center for Brain Imaging cost researchers around $450 per hour of subject time. In addition to the price, collecting neuroimaging data can have many other non-monetary costs such as requiring the subject to come into the lab, requiring the time of trained researchers or technicians to run the machine and only being able to run one subject at a time. In contrast, many behavioral studies can be easily run in parallel online without significant monitoring by researchers for a cost of around $11 per hour of subject time (Crump et al., 2013; Hara et al., 2018). Thus, if 40 behavioral subjects can improve inferences more than a single fMRI subject, this might be a worthwhile tradeoff in many experimental designs.

To make our results more general, we compare estimates using five values of *N*_*xy*_ (25, 50, 100, 500 and 1000) and five values of *ρ* from .1 to .9. For each set of (*N*_*xy*_, *N*_*x*_, *ρ*), we generate 1,000,000 datasets and estimate the correlation using both methods. In Fig. 5, we plot the benefit of collecting additional behavioral data in terms of the change in root mean squared error of collecting an additional neuroimaging subject and using the standard Pearson *r*. The grey dotted lines at 0 and −1 in Fig. 5 reflect the blue and orange lines in Fig. 4. The solid lines indicate the performance of the Anderson (1957) estimator (the green line in Fig. 4). In practice, the Anderson (1957) estimator is not beneficial in all conditions. This is indicated by the color of the solid line: in cases where it does not benefit, we color the solid line red.

**Figure 5:**
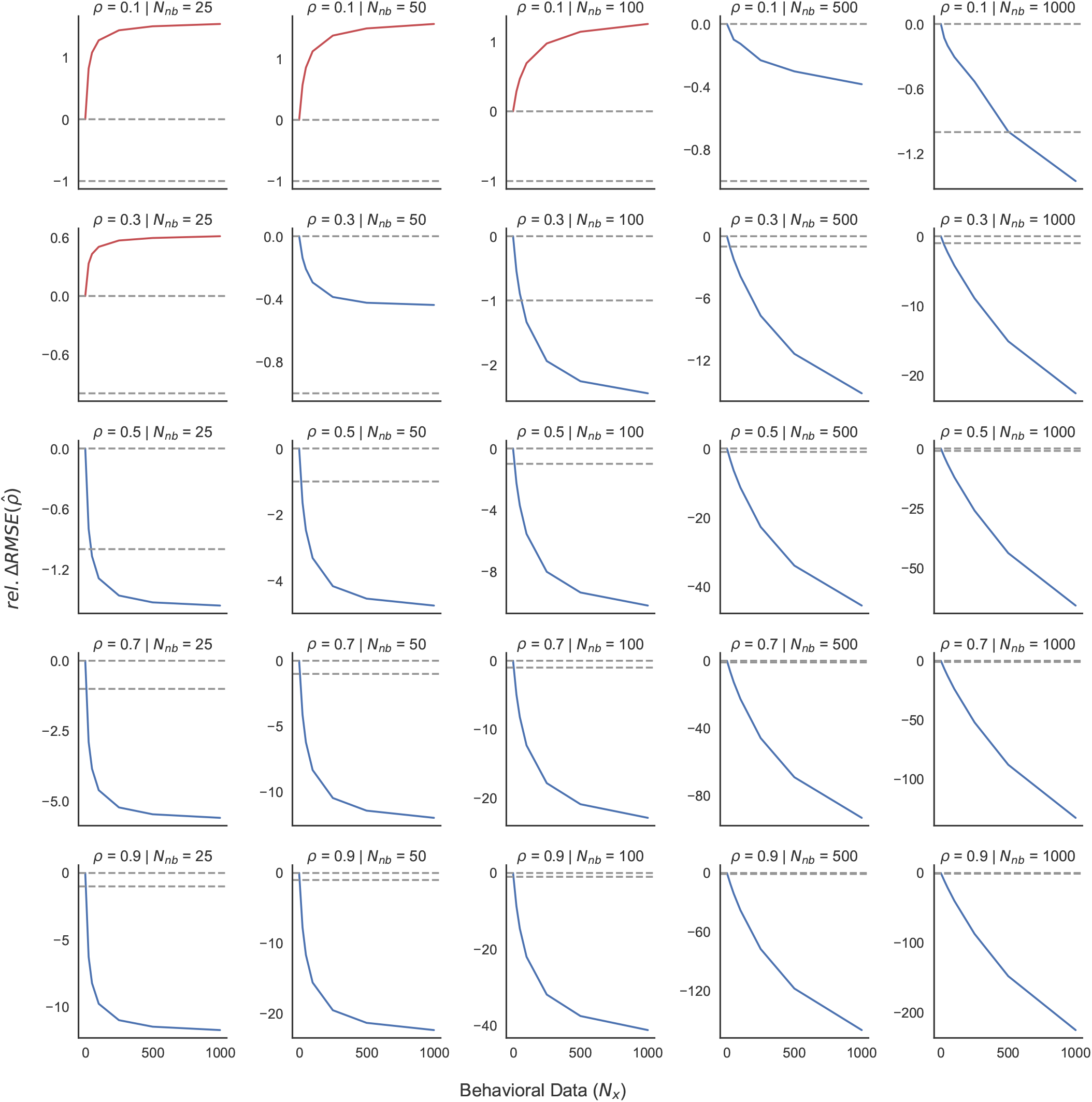
Relative decrease in root mean squared estimation error, averaged over 1,000,000 simulated datasets, for the estimate of *ρ* from Anderson (1957) as a function of the true correlation, the number of neuroimaging subjects *N*_*xy*_ and number of additional behavioral subjects *N*_*x*_. Values are computed relative to the change in RMSE for a Pearson correlation with *N*_*xy*_ neuroimaging participants (0) compared to *N*_*xy*_ + 1 neuroimaging participants (−1). These values are indicated by the grey dashed lines. The solid line shows the Anderson (1957) 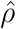 changes in the estimate as the number of addition behavior only subject are added to the analysis. The solid lines are colored red in the cconditions where the Anderson (1957) estimate is not beneficial and blue in the majority of cases where it is. Note that the scale of the y-axes are not fixed across plots but the grey dashed lines have the same meaning across plots. This is necessary in order to see the change with number of subjects.

Because we are emphasizing the tradeoff between collecting neuroimaging and behavioral data, the y-axis of Fig. 5 is plotted relative to the precision benefit of one additional neuroimaging subject. However, in the supplemental material (Fig. 10), we also show the absolute change in precision (in terms of root mean squared error) as a function of the parameters we discuss here.

Comparing the two methods of correlation estimation, we find that the effect of the additional behavioral data can be nonlinear in the size of the underlying neuroimaging dataset and the true value of *ρ*. Having too few neuroimaging data points to estimate a weak correlation (e.g. only 25 subjects for a correlation of .1) makes the behavioral data less useful and it can be better to collect more neuroimaging data. This finding resembles similar simulation results from Garren (1998). At low amounts of neuroimaging data and lower true values of *ρ*, the Anderson estimate of the correlation using all of the behavioral data can actually perform worse than the standard Pearson estimate. However, as the true *ρ* and number of neuroimaging subjects increase, the value of the additional data increases such that collecting additional behavioral data can be better than collecting a smaller amount of additional neuroimaging data. In some cases, collecting just 25 behavioral subjects can improve precision more than another neuroimaging subject. At the cost levels described above, this can mean that behavioral data can be a better investment than equivalently priced neuroimaging data. Therefore, a design analysis that takes costs into account has the potential to recommend collecting behavioral data rather than neuroimaging data.

On the other hand, we reiterate that this is not a general recommendation. As shown in Figure 5, it is not true across the whole parameter range and, perhaps in particular, it isn’t true for low sample sizes and low true correlation values, which is exactly where we might hope to gain in terms of precision. Why is it that the Anderson estimator performs worse here?

To answer this question, we can decompose the mean squared error into the bias and the variance, i.e.

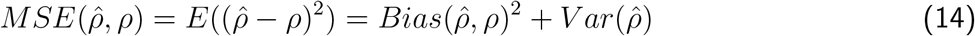

where Bias is defined as

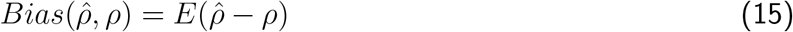

As with mean squared error, we can also look at how various methods perform in terms of bias, i.e. is the estimate of *ρ* equal to *ρ* on average? It is well known that Pearson’s *r* is biased downward such that, on average, correlation estimates will be lower than their true value (Olkin & Pratt, 1958). In Figure 6, we can see that the Anderson estimator of *ρ* always decreases the bias in estimating *ρ*, relative to an *N*_*x*_ of 0 where it is equivalent to the Pearson *r*. This means that the increase in mean squared error in the Anderson estimate for lower values of *ρ* and *N*_*xy*_ is due to the behavioral data adding variance to the estimate. It may be that the adjustment shown in Figure 3 is only likely to be in the right direction when there is enough data to estimate *ρ* and *σ*_*x*_ precisely. We hope that future statistical work will address this question and potentially develop new estimators that can control this variance better in the low sample size, low correlation regime. For now, the choice of whether to use the Anderson (1957) estimator will depend on the research question. In the appendix, we investigate the performance of the Anderson estimator in real data from the Human Connectome Project (Van Essen et al., 2013) and show that it largely matches the performance expected from these simulations (Fig. 11).

**Figure 6:**
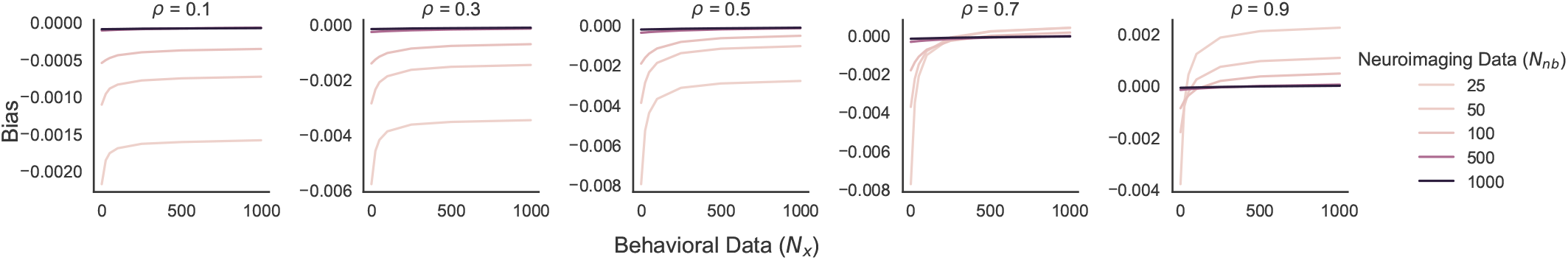
Bias in the Anderson (1957) estimate of *ρ*, over 1,000,000 simulated datasets, as a function of as a function of the true correlation, the number of neuroimaging subjects *N*_*xy*_ and number of additional behavioral subjects *N*_*x*_. Note, as in the previous figure, that the axes here are not fixed across plots. This is necessary in order to see the change with number of subjects.

A side benefit of the above derivation in Eq. 15 is that it is easy to see how the 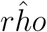 estimate depends directly on variance estimates. At least since Gauss, statisticians have known to include an adjustment for the degrees of freedom when estimating an unbiased variance from a sample. In the traditional Pearson estimate of the correlation, these adjustments do not appear since they cancel out when the covariance is divided by the variance. However, this is not true of the Anderson *ρ* estimator, due to the 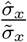 term. In the original formulation Anderson (1957) and in all other treatments we have seen, researchers use the biased maximum likelihood estimate. However, we find that adjusting for the degree of freedom does improve estimates in the more likely scenarios of low numbers of joint data and low true correlations (Fig. 12). We hope that future work can build on these results to discover improved estimators.

One somewhat counter-intuitive consequence of the above discussion is that Anderson’s estimator can help not only in studies of individual differences across subjects but also in studies using representational similarity analyses where the matrix of item-item similarities based on a model is correlated with the similarities based on the multivariate neural signal (Edelman et al., 1998; Kriegeskorte, 2008). If researchers simply increase the number of similarities computed for the model, the above analyses suggest that there may be cases where this increases the power to detect a correlation with neural measures. In practice, since representational similarity matrices are often created from behavioral ratings (e.g Bruffaerts et al., 2013; Chikazoe et al., 2014), this is yet another way in which collecting additional behavioral data can increase power. While it is common in this literature to use Spearman’s rank correlation to assess similarities, we can put this within the linear correlation framework by first ranking the two variables and then computing the correlation. For Pearson’s *r*, this is exactly equivalent to computing the Spearman correlation.

### 3.2 How to Decrease Estimation Error: Improving Within-Subect Model Estimates by Collecting Data from Additional Subjects

We now investigate how collecting additional behavioral data can also help reduce estimation error, making it applicable to the regression analyses described above. In our previous description of estimation error, we were agnostic as to how exactly parameters were estimated from behavioral data. The most common method of estimating cognitive model parameters for use in a **model-based regression** is to use maximum likelihood estimation (Myung, 2003). The maximum likelihood estimate of parameters *θ*_*i*_ for subject *i* is the set of parameters in a cognitive model *C* that maximizes the probability of that subject’s behavioral data **x**_*i*_ i.e.

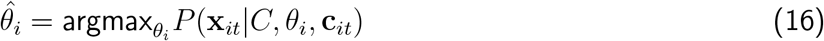

One problem is that many neuroimaging experiments are relatively short so not many trials can be collected per subject. In complex cognitive models, when the number of data points is small, many sets of parameters can be consistent with the data and, therefore, two datasets generated from the cognitive model with the same *θ*_*i*_ could result in very different maximum likelihood estimates 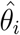, i.e. 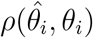 can be low on average. As demonstrated in the above estimation error section, this will result in lower power for estimating correlations and regression slopes when attempting to relate such variables to neural recordings.

Two methods have been used in the literature to increase 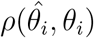 and 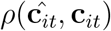: sharing information across subjects or constraining the possible parameter values. One way to use information from other subjects data is to simply assume that all subjects have the same *θ* values which means that they can be estimated using all of the behavioral data collected in the experiment (Daw, 2011). This increases the data used to fit the parameters by a factor of *N*_*xy*_. The **c**_*it*_ will still vary by subject because of the dependence on the stimuli in the experiment. If the variability in true parameters *θ*_*i*_ across subjects is small relative to the estimation error, this can be an effective, heuristic way to increase *ρ*(**ĉ**_*it*_, **c**_*it*_). However, this is not an option in a **correlation** analysis when we are often interested the variation across subjects itself. In addition, many cognitive processes do have large individual differences so this may actually decrease *ρ*(**ĉ**_*it*_, **c**_*it*_).

Another option is to constrain parameter values by adding a prior, *P* (*θ*_*i*_|*ϕ*). If *ϕ* is chosen correctly, using a prior can decrease the likelihood of implausible parameter values such as a learning rate of 0 (Daw, 2011). We can then find the parameters that maximize the posterior, i.e.

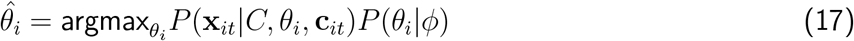

This is known as the maximum a posteriori (or MAP) estimation. By using the prior, we can force *θ*_*i*_ to be closer to more plausible values. However, it can be difficult to know in general if your priors are constraining the parameters to be in the correct space. If your priors are wrong, this could also decrease 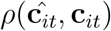.

One principled way to incorporate information from other subjects and use priors is hierarchical modeling (Gelman, 2006). This is an increasingly popular method in cognitive science and cognitive neuroscience (Ahn et al., 2011; Rouder & Lu, 2005) that assumes that there is a potentially nonuniform distribution *P* (*θ*_*i*_|*ϕ*) of parameters in the population. Functionally, this constrains the possible values of *θ*_*i*_ as in the prior above but because this is describing the population distribution, we can now estimate *ϕ* from all of the subjects data rather than setting *ϕ* by hand. The resulting estimate of *θ*_*i*_ will be a weighted average of the estimate from the population with the estimate that just uses an individual’s data. The weights are determined by the strength of the individual data. For a subject whose data provides less constraint on the parameters of the cognitive model (i.e. if they had fewer trials or made more inconsistent choices), their estimates will be closer to the prediction from the prior. If a subject made very consistent choices, we might already have a lot of information about the parameters of their cognitive processes and we do not need to use as much information from the population. Thus, hierarchical modeling provides a way to decide from the data how much we should pool information from the population in creating each individual estimate. Katahira (2016) demonstrates that estimates from a hierarchical model strictly dominate both individual maximum likelihood estimates and population maximum likelihood estimates in terms of bias and variance.

While there exist frequentist methods for estimating hierarchical models (such as restricted maximum likelihood as implemented in popular packages for fitting hierarchical linear models like `lmer` (Bates et al., 2015)), for arbitrary cognitive models, fitting in this way often requires a complicated derivation or numerical integration which can be challenging in high dimensions. With modern Bayesian inference tools like `Stan` (Carpenter et al., 2017), it is usually much more straightforward to place a prior on *ϕ* and use Bayesian inference to compute a joint posterior over *ϕ* and *θ*. The posterior for the hierarchical model is then

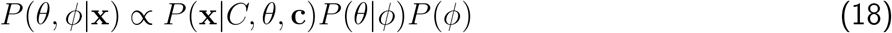

In addition to Stan, many R or python packages have made hierarchical Bayesian versions of specific cognitive models particularly easy to fit (e.g. (`hddm` (Wiecki et al., 2013) for drift diffusion models or `hBayesDM` (Ahn et al., 2017)) for many types of decision making models).

One feature of hierarchical modeling is that individual hierarchical estimates can often be improved simply by sampling more data from the population. That is, the cognitive model parameter estimates for subjects whose data you already have, i.e. subjects who contributed neuroimaging data, can be improved by adding data from new subjects. This is because the error of the estimates of *ϕ* (or the standard deviation of the prior) will, for most models, converge towards the correct estimates with increasing the number of subjects.

To demonstrate this, we conduct a simulation with a very simple hierarchical model to show that additional behavioral data from new subjects can improve inference for latent cognitive parameters *θ*_*i*_ for subjects *i* ∈ [1, (*N*_*xy*_ + *N*_*x*_)], *N*_*xy*_ of whom were collected in an original set neuroimaging experiment and *N*_*x*_ who were collected in a separate behavioral experiment. We assume each subject’s latent cognitive parameter *θ*_*i*_ is drawn from a Gaussian population distribution with mean *µ*_*θ*_ = 0 and variance 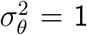. Each subject then provides behavioral data *b*_*it*_ and to keep things simple, we assume that the cognitive state variables *c*_*it*_ are essentially equal to *b*_*it*_ such that

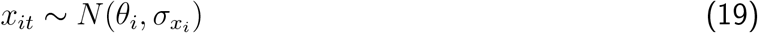

where *σ*_**x***i*_ is drawn from a uniform distribution between .5 and 1 for each subject. Further simplifying, we summarise *x*_*it*_ by its mean 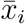 and assume that there are enough trials *t* that we can treat 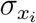 as known.

For the hierarchical model, we need to put a prior on 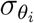. While in practice it is often better to use a weakly or strongly informative prior, for generality, in these simulations, we assume an improper uniform prior, i.e. *P* (*σ*_1_) ∝ 1. Using the derivation in (Gelman et al., 2013), we construct a grid approximation to the posterior for *θ* given **x**. We now investigate how the correlation between the true subject parameters *θ*_*i*_ and their posterior mean estimate 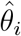 for the *N*_*xy*_ neuroimaging subjects changes as we add a number of new behavior-only subjects, *N*_*x*_, to the experiment. Note that we are only computing the correlation within the original *N*_*xy*_ subjects. As we demonstrated in the section on estimation error, this correlation is what limits the power to detect true correlations or regression slopes between model estimates and neural signals.

Figure 7 shows that across a range of *N*_*xy*_ values, the addition of *N*_*x*_ new behavioral subjects can increase the correlation between model estimates for the *N*_*xy*_ neuroimaging subjects 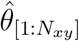 and true parameters 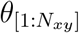.

**Figure 7:**
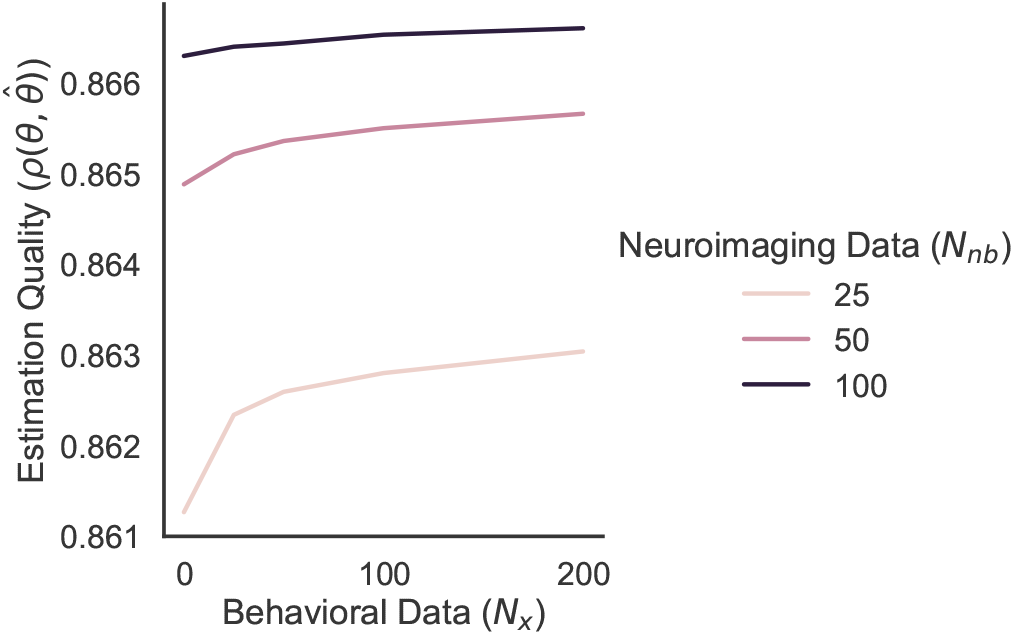
Correlation between the estimated cognitive trait parameters 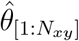 and 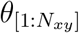 among the *N*_*xy*_ neuroimaging participants using the hierarchical model, averaged over 10,000 simulated datasets, as a function of the number of neuroimaging subjects *N*_*xy*_ and number of additional behavioral subjects *N*_*x*_.

This implies that one way to increase 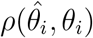 (and therefore also increase *ρ*(**c**_**i**_, **ĉ**_**i**_)) for subjects in a neuroimaging experiment is to simply collect behavioral data from more subjects. However, the exact rate at which more subjects improve the model estimates will in general depend on the distribution of uncertainty in the individual *θ*_*i*_ as well as the particular form of the model you use. We also emphasize that while the increases in 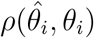 may be modest, for particular cognitive models, a small improvement in trait estimates could lead to a large improvement in state estimates *ρ*(**c**_**i**_, **ĉ**_**i**_). It is these state estimates that are most commonly of interest in a model-based neurogimaging analysis (e.g. reward prediction error in a reinforcement learning model).

Beyond its use in improving hierarchical estimates of cognitive model parameters, collecting additional behavioral data can improve neuroimaging regression inferences in several other ways. While not computationally feasible yet for most neuroimaging analyses, joint modeling (B. M. Turner et al., 2013) is a recently proposed technique in which the cognitive model and the neuroimaging model are jointly fit, hierarchically, to both the neuroimaging and behavioral data. This can allow for even more efficient use of information by allowing both sources of data to inform the cognitive model parameters. In addition, it allows the uncertainty in the cognitive model parameters to be propagated into the estimates of the regression or correlation, resulting in more powerful estimates without the biases mentioned above. B. M. Turner et al., 2016 demonstrated how joint modeling can be used to improve multimodal inferences even when we don’t have data for every mode for every row in the dataset (i.e. some subjects’ data were collected with EEG and others with fMRI). This can be generalized to the case we discuss where we only have rows in the dataset with only one type of data (e.g. behavior). Collecting additional behavioral data in experimental conditions or on novel stimuli not in the neuroimaging experiment could additionally assist in model selection. Choosing the correct model could potentially have a much larger impact on the correlation between model estimates and true parameters.

Finally, we can view reducing estimation error as relevant not only for model-based regression but also for regression approaches where the stimulus features are subjective or latent, such as the valence of a stimulus. The covariates in this case are often created from ratings based either on subjects in the experiment or outside behavioral subjects. Similar to their use with cognitive models, hierarchical models with additional behavioral data can reduce error in these stimulus ratings as well. Even just aggregating the average rating however, will be improved through greater amounts of data per item. Indeed, going back to at least Kent and Rosanoff (1910), it is already quite common in cases like this to use large “norming” studies to understand stimuli, particularly for word stimuli.

#### 3.2.1 When does it work?

Having demonstrated that it is possible to reduce estimation error in a neuroimaging dataset by collecting more behavioral data, we might now wonder when this is likely to help. As we mentioned above, the amount that hierarchical modeling can help will in general depend on the model and the experimental design. In addition, collecting additional behavioral data can improve estimates in other ways such as through model selection. In order to say something general about how improvements in estimation improve statistical power to find neuroimaging effects, we will simply assume that it is possible to increase the correlation between model estimates and true parameters using behavioral data by a certain amount and investigate how that changes estimates about the relationship with neural data.

In this paper, we have assumed that collecting data from additional subjects is the cheapest way to decrease estimation error (due to the availability of easy online data collection). In addition, we argued above that collecting data from more subjects reduces estimation error in hierarchical model estimates in a straightforward way through improved hyperparameter estimation. However, collecting additional behavioral data from the *N*_*xy*_ neuroimaging subjects will, in many cases, also increase 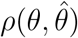. For instance, increasing the number of trials will necessarily decrease the standard error of the mean of some quantity computed over all of those trials. Which strategy is better will depend on many details but, as our analysis is in terms of the increased correlation between model estimates and true parameters rather than in terms of number of subjects, our results will apply equally to both strategies of reducing estimation error.

To begin, we will build on the derivation from Wilson and Niv (2015) where they showed how the least squares estimate of the regression slope is impacted by misestimation (i.e., equations 4 and 5). Wilson and Niv (2015) only solved this for the case where the analyst is trying to infer a single regression slope from the data (i.e. a non-hierarchical regression model). But in model-based neuroimaging analyses, it is rarely assumed that all subjects have the same effect size. Therefore, the most common analysis method in model-based analysis is to use a two stage approach where a first-stage regression is fit to each subject (or each run) and then the subject regressions are aggregated at the group-level by approximating a hierarchical model, testing whether the average effect is significantly different from zero (Beckmann et al., 2003; Woolrich et al., 2004). This means that the benefit of collecting an additional neuroimaging subject cannot be captured by the Wilson and Niv (2015) equations alone. Following Friston et al. (2002) and Beckmann et al. (2003), we assume a Gaussian hierarchical model of population effects:

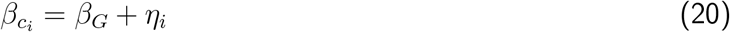

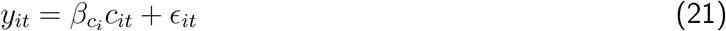

where the noise distributions are both Guassian:

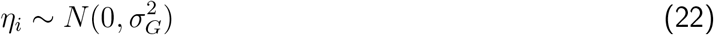

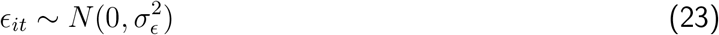

Using the same notation as above, for subject *i* on trial *t, c*_*it*_ is that subject’s cognitive state at that time and *y*_*it*_ is the associated neural signal. 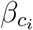 represents the relationship between the cognitive state and the neural signal for a particular subject and *β* represents that average relationship in the population. 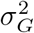 represents the variance of the magnitude of the relationship in the population and 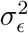 represents the additional variance in the neural signal that is not due to the cognitive state (either due to noise in the measurement or the signal being related to other unrelated cognitive processes as well.)

In order to quantify the value of the improved model fit, we first must derive the power for estimating a reliable *β*_*G*_ parameter given a particular relationship between the true model parameters and the model estimates. In the Appendix, we obtain an analytic expression under the assumption that the second-level model is estimated using the standard OLS method.

Rather than parameterizing this in terms of the raw effect size, *β*_*G*_, it turns out to be more illuminating standardize the variables by the total standard deviation, 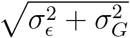, and, following Wilson and Niv (2015), we call this standardized effect size the contrast-to-noise ratio (CNR). Additionally, in multilevel modeling, an important statistic is the intra-class correlation coefficient or ICC which is the proportion of the total variance that is explained by the subject level variance, i.e. 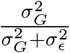 (Chen et al., 2018; Shrout & Fleiss, 1979). Given these terms, we get that the expected *t*-statistic under a particular study design is

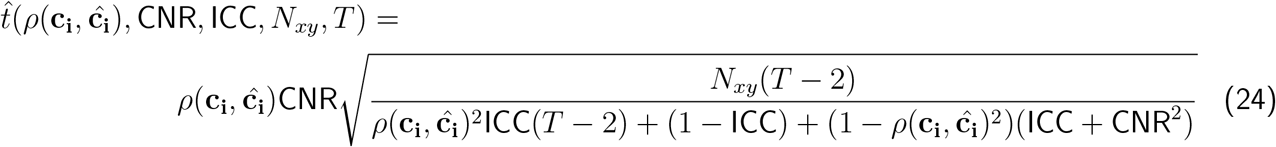

We can then compute the power using standard formulas and in Figure 8, we show how it depends on the model fit *ρ*(**c**_**i**_, **ĉ**_**i**_) for several values of CNR, *T* and *N*_*xy*_. As expected, improving the model fit increases power, although in a highly non-linear way, meaning that the tradeoff for improving model-fit depends on all of these parameters.

**Figure 8:**
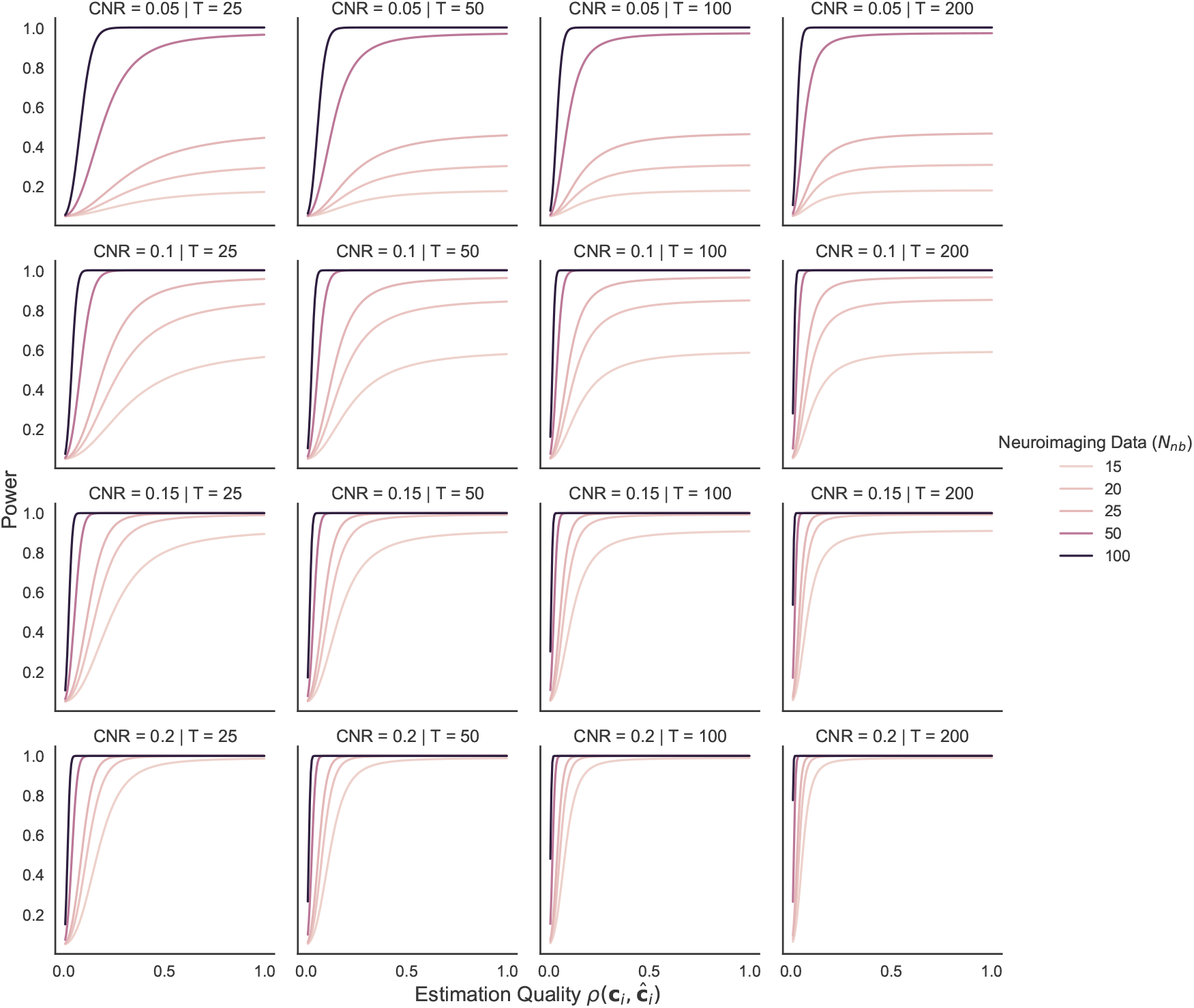
Power as a function of *ρ*(**c**_**i**_, **ĉ**_**i**_) for a range of CNR, *T* and *N*_*xy*_ values. ICC was set to .397, corresponding to a recent estimate in the literature (Elliott et al., 2020). In general, power increases as the number of subjects increases, as the number of trials increases and as accuracy of the model estimates (the inverse of the estimation error) increases.

This leads us to ask the question “how much would additional behavioral data have to improve our latent variable estimates in order to improve power as much as another neuroimaging subject?” That is, what is the increase (δ_*ρ*_) in the correlation between estimated model parameters and true parameters (*ρ*(**c**_**i**_, **ĉ**_**i**_)) that is equivalent to increasing *N*_*xy*_ by one? More formally, if *P* (*t, N*) is the power of the one-sample *t*-test with a *t*-stat of *t* and *N* subjects, we can write

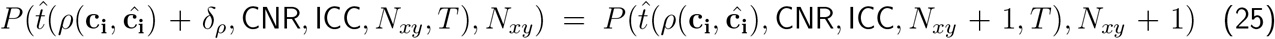

and ask what value of δ_*ρ*_ makes this equation true?

In order to keep our results general, we analyze this situation in terms of how much the correlation would have to increase, a dimensionless quantity, rather than a number of data points. This is because, because, as we noted above, this amount will depend on the specific cognitive model. In order to translate (δ_*ρ*_) into a specific amount of subjects, one could conduct a simulation similar to the one displayed in Fig. 7 to obtain the relationship between *N*_*x*_ and *ρ*(**c**_**i**_, **ĉ**_**i**_) for the specific cognitive model of interest.

Since δ_*ρ*_ must be between 0 and 1 − *ρ*(**c**_**i**_, **ĉ**_**i**_), this is a straightforward constrained root finding problem that we can solve exactly using numerical methods. We can solve this for several values of the other parameters (CNR, ICC, *T* and *N*_*xy*_) using Brent’s method (Brent, 1972) as implemented in the `scipy` package (SciPy 1.0 Contributors et al., 2020) in `python 3.7.7`.

Which settings of the parameters in equation 25 are reasonable for neuroimaging experiments? Wilson and Niv (2015) derive estimates for CNR from two different studies with a very wide range of .4 to 11. Their CNR definition was only normalized by *σ*_*ϵ*_ and not the total standard deviation including the group variance as we have done here. Therefore, to translate those values to values in equation, we need to take into account the ICC. A recent meta-analysis of reliability across a wide-range of task-based neuroimaging studies found an average ICC of .397 (Elliott et al., 2020). With that value of ICC, Wilson and Niv (2015)’s CNR of .4 corresponds to a CNR of 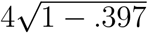 which is approximately .31. We can see in Figure 8 that a value of even .2 for the CNR leads to somewhat implausible power estimates for most cognitive neuroscience task-based studies (Button et al., 2013; B. O. Turner et al., 2018). Therefore, we will assume that the CNR is closer to .1 in the following simulation. However, with the above derivation, it is easy to compute this for other values of the generative model and task parameters. Also of note in these plots is that for a particular value of *N*_*xy*_ the power function will asymptote as a function of *ρ*(**c**_**i**_, **ĉ**_**i**_). This means that given a good enough model estimate, it is impossible to improve power more by improving the estimate than by collecting more subjects.

Figure 9 shows the solution to equation 25 in terms of δ_*ρ*_ (the improvement to match the estimate with an additional neuroimaging subject) as a function of the correlation between estimate and true parameters with one more neuroimaging subject. Overall, as noted above, there is a range of parameter settings where improving model fit (potentially by collecting more behavioral subjects) cannot help improve the power more than collecting another neuroimaging subject. Wilson and Niv, 2015 showed that, for a reinforcement learning model, it is fairly easy to obtain a high *ρ*(**c**_**i**_, **ĉ**_**i**_), even if the latent subject-level parameters are severely misestimated. This suggests that in that setting, model-fitting “isn’t necessary”, or at least improving the model fit may be unlikely to significantly affect power. However, this may not be the case for other cognitive models, although it seems unlikely that a model that is fit to the data will have a correlation as low as .2 or .3.

**Figure 9:**
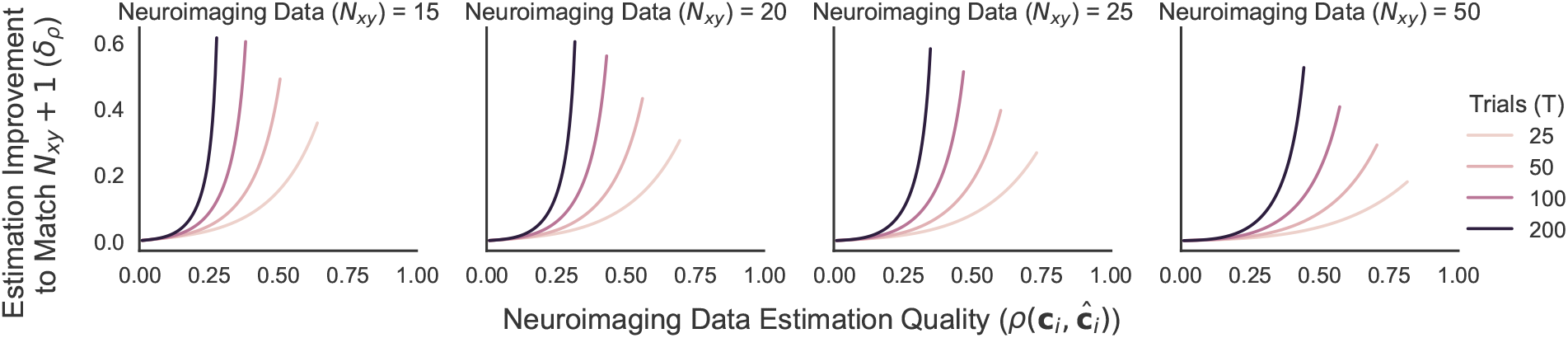
Solution to equation 25, showing the needed improvement in estimation quality to match an additional neuroimaging subject as a function of parameters *ρ*(**c**_**i**_, **ĉ**_**i**_) for a range of the number of trials (*T*) and the number of neuroimaging subjects (*N*_*xy*_). CNR is set to .1 and ICC is set to .397 (Elliott et al., 2020). Lines end when it is no longer possible to achieve better performance by increasing the correlation.

In future work, we hope to explore this in the context of other cognitive models. Assuming that correlations between are reasonably high and that a smaller δ_*ρ*_ is easier to achieve, Figure 9 shows that improving the model fit with additional behavioral data is most useful when there is already a reasonably sized *N*_*xy*_ and *T* is smaller. While even 50 subjects is already significantly above average for a neuroimaging study, most use numbers of trials that are much higher than 50. One thing that this suggests is that using extra behavioral data may be particularly useful for neuroimaging studies with small numbers of trials. These are not common because of the low power they achieve but methods like those mentioned in the previous section may make them more feasible. This opens the door to more neuroimaging studies that are difficult to run with high numbers of trials such as long-term memory studies where subjects are unlikely to be able to remember a large number of items over a long delay. Finally, in this section, we focus on inferring the conditional relationship between cognitive model parameters and neural signals, assuming that the linear model is correct. However, there are many situations where it might be statistically reasonable or even preferable to use a linear model even if it is known that the neural signals are not exactly a linear function of the cognitive parameters (Buja et al., 2019; White, 1980). Recent work by Azriel et al. (2021) has shown that in this case, inference for regression parameters can be improved by additional unlabeled data as we show in the above work. Future work should also explore the value of behavioral data for model-based fMRI in this approximation regime.

## 4 Discussion

In this paper, we have presented two ways in which collecting additional behavioral data outside of a neuroimaging experiment has the potential to improve the precision of an estimate of the relationship between brain measures and cognition. We also attempted to quantify a tradeoff between collecting each type of data, showing how this could potentially affect the design of future studies or reanalyses of past work. In both cases, behavioral data had the potential to help but not in all cases.

Critically, our simulations reveal specific parameter regimes where these techniques are most effective, serving as a guide for researchers considering this approach. Regarding sampling error, we found that the benefit of the Anderson estimator is non-monotonic; it provides the substantial gains over collecting additional neuroimaging data when the original sample size is moderate (e.g., *N >* 25) and the true underlying correlation is moderate-to-strong (*ρ >* 0.3). Conversely, in very small samples looking for weak effects, the added variance from the behavioral data can outweigh the benefits. We hope that these somewhat mixed results can spur future work on how to most effectively leverage missing data approaches to combining neural and behavioral data for estimating correlations.

In contrast, the benefit of reducing estimation error via additional behavioral subjects (e.g., through hierarchical modeling) follows a different pattern. This approach is most advantageous when the precision of individual subject estimates is the limiting factor—specifically in studies with low trial counts (*T*) or high measurement noise. Thus the hierarchical approach provides a clear benefit allows researchers to maximize power in resource-constrained neuroimaging designs where collecting many trials per subject is infeasible.

The ideas proposed here do have additional limitations as well. For instance, while in this paper we have assumed that data collected with and without a neuroimaging recording device (such as in an fMRI scanner or with an EEG cap on) are exchangeable, that is unlikely to be exactly true in practice. This does not make the described methods impossible to use but does potentially require more complex models that allow for differences between the data collected under different modalities. That being said, there is relatively little work documenting the magnitude of these purported differences. If these differences were found to be large, this would likely have implications for learning about cognition through neuroimaging in general, not just with the statistical methods described here. We strongly encourage both future empirical work documenting these differences and future statistical work on how to aggregate inferences across data collection modalities.

Our analyses suggest important directions for future statistical work, in particular the potential to improve the Anderson (1957) correlation estimate by appropriately reducing the variance, especially in situations with low true correlations. In addition, our analyses suggest new regimes where behavioral data could be helpful; for instance, the potential for improving model fits with behavioral data could allow for feasibly running neuroimaging studies with small numbers of trials. In many cases, such as in long term memory studies, it is not feasible to run large numbers of trials because participants cannot reasonably remember hundreds of items. Both of these directions will require more work to be put into practice but we hope this paper will inspire collaborations between cognitive neuroscientists and statisticians to work more on ways of fusing data sources in order to learn about brain and behavior relationships.

We are, of course, not the first to have pointed out that collecting data with some modalities missing can still be used to improve inferences. Indeed, this was the perspective taken in B. M. Turner et al. (2016). We believe that we advance the literature in several important ways: 1) we connect this idea to the existing statistical literature with missing data; 2) We show that we can improve inferences not only for parameters of a cognitive model but also for the estimates of the relationship between brain and cognition, often a more important target of inference in cognitive neuroscience; 3) we provide new derivations that help understand (and improve, Fig. 12) the Anderson *ρ* estimator, which we believe is a starting off point for combining behavioral data with neuroimaging data; 4) we provide a general treatment of the issue which is not tied to a particular cognitive model or dataset.

**Figure 10:**
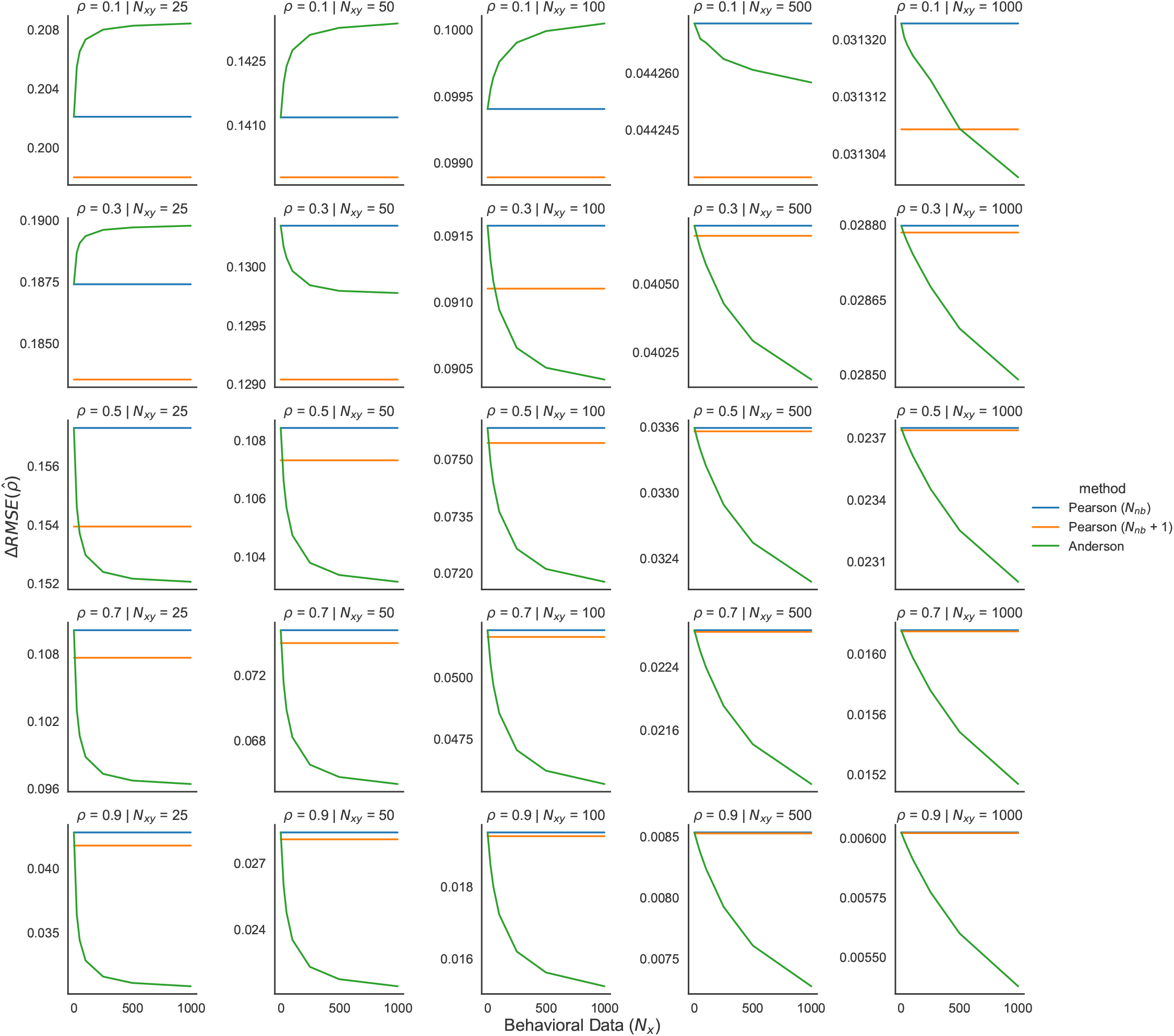
Decrease in root mean squared estimation error, averaged over 1,000,000 simulated datasets, for the estimate of *ρ* from Anderson (1957) as a function of the true correlation, the number of neuroimaging subjects *N*_*xy*_ and number of additional behavioral subjects *N*_*x*_. As in Fig. 4, we also plot Pearson correlation with *N*_*xy*_ neuroimaging participants and *N*_*xy*_ +1 neuroimaging participants, indicated by the orange and blue lines. The green line shows the Anderson (1957) 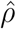 changes in the estimate as the number of addition behavior only subject are added to the analysis. Note that the scale of the axes here are not fixed across plots. This is necessary in order to see the change with number of subjects.

**Figure 11:**
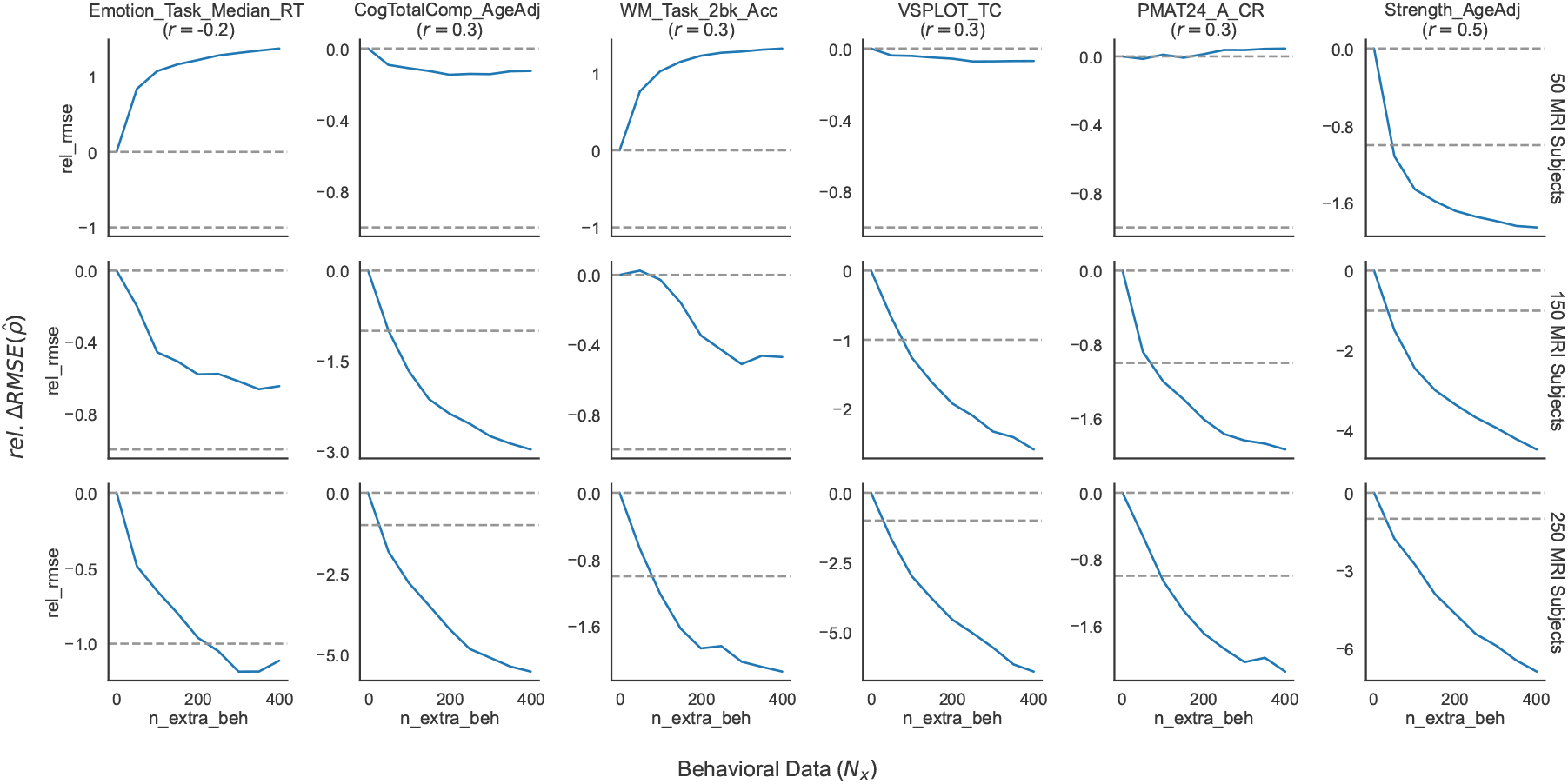
Relative decrease in root mean squared estimation error, averaged over 50,000 boothstrapped datasets, for the estimate of *ρ* as a function of the variable (with the correlation in the full dataset indicated in parentheses), the number of neuroimaging subjects *N*_*xy*_ and number of additional behavioral subjects *N*_*x*_. Values are computed relative to the change in RMSE for a Pearson correlation with *N*_*xy*_ neuroimaging participants (0) compared to *N*_*xy*_ + 1 neuroimaging participants (−1). These values are indicated by the grey dashed lines. The solid line shows the Anderson (1957) 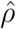 changes in the estimate as the number of addition behavior only subject are added to the analysis. Note that the axes here are not fixed across plots. This is necessary in order to see the change with number of subjects.

**Figure 12:**
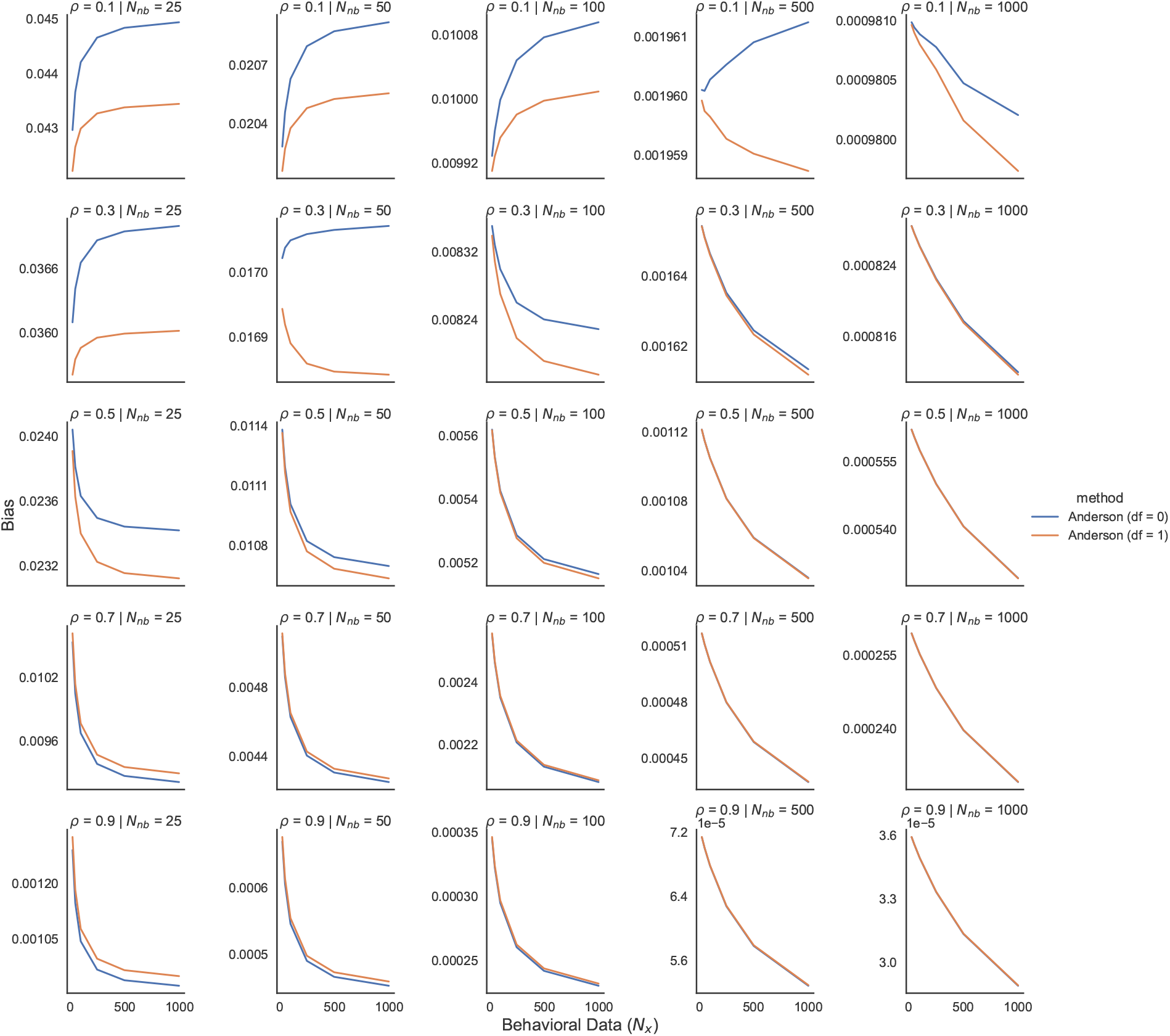
Root mean squared estimation error, averaged over 1,000,000 simulated datasets, for the estimate of *ρ* using the unbiased (df = 1) and biased (df = 0) estimates of variance, as a function of the true correlation, the number of neuroimaging subjects *N*_*xy*_ and number of additional behavioral subjects *N*_*x*_. Note that the axes here are not fixed across plots. This is necessary in order to see the change with number of subjects.

While we have mostly discussed collecting additional behavioral data in the context of designing a new study, it could also be a consideration in a study that resuses an old dataset. It could be that a model is not identifiable with a particular dataset but it could become identifiable by collecting behavioral data in more conditions. With the advent of more open neuroimaging data sets, and particularly with the current difficulty in collecting new data, these methods point to a way to learn new things from old datasets simply by collecting more cheap behavioral data.

## Data and Code Availability

Code will be available upon acceptance for publication. There is no data associated with this paper.

## Author Contributions

David Halpern: Conceptualization, Methodology, Investigation, Formal analysis, Visualization, Writing—Original draft. Todd Gureckis: Conceptualization, Writing—Review & editing, funding acquisition.

## Declaration of Competing Interests

The authors declare that they have no competing interests.

## A Appendix

### A.1 Derivation of relationship between Pearson *r* and Anderson 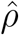

Here, we try to illuminate how the new estimate is different from the correlation estimate based on only the *N*_*xy*_ complete cases. Little and Rubin (2019) derive the missing data estimator 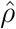 in a different way, showing that

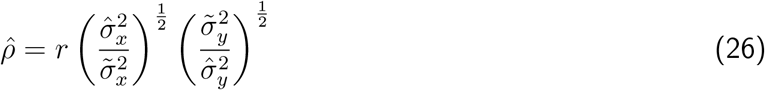

where *r*, 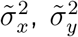 are the estimates of the correlation (i.e. Pearson’s *r*), the variance of the behavioral variable and the variance of the neural variable among the *N*_*xy*_ complete cases. 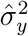 is the variance of *y* after adjusting for the 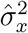estimate based on all of the data. Little and Rubin (2019) derive the formula for 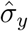 as

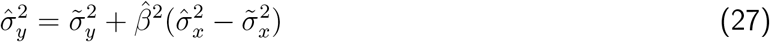

We can instead write this in terms of *r* and rearrange terms, i.e.

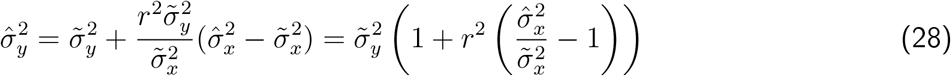

Plugging this back in, we get

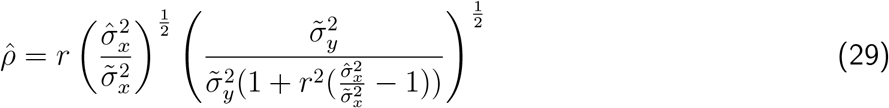

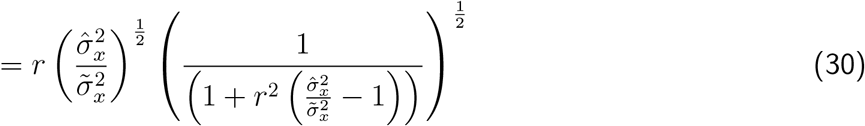

### A.2 Absolute Change in Precision for Anderson’s 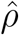

While in the main text, we emphasize the benefit of behavioral data *relative* to an additional neuroimaging subject, it is also of interest to know the absolute change in precision when using the Anderson estimator. Therefore, we re-plot 5 in terms of root meaa square error.

### A.3 Application to Human Connectome Data

In order to test the performance of the Anderson 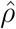 estimator, we conducted an analysis of brain-behavior correlations in the Human Connectome project (Van Essen et al., 2013) public data. Specifically, we investigated the relationship between total grey matter (FS_Total_GM_Vol) and performance on several cognitive tasks: The Variable Short Penn Line Orientation (VSPLOT_TC) (Benton et al., 1975; Gur, 2001), a two-back working memory task (WM_Task_2bk_Acc) (Cohen et al., 1994; Kirchner, 1958), the Penn Progressive Matrix Task (Gur, 2001; RAVEN, 1941), a Language comprehension task (Language_Task_Acc) (Binder et al., 2011) and the age-adjusted composite Cognitive Function score from the NIH Cognitive Toolbox (CogTotalComp_AgeAdj) (Heaton et al., 2014). In order to generalize to a wider variety of task types and measures, we also included response times in a social-cognitive emotion task (Hariri et al., 2002) and grip strength (Reuben et al., 2013). In the full dataset of 1076 subjects with data on all variables, these correlations ranged from .2 to .5. We created 50,000 bootstrapped datasets and analyzed the variance of estimated correlations using Pearson’s *r* and the Anderson *ρ* estimator, assuming that the correlation on the full dataset was equal to the “true” population value (Fig. 11). As expected from the simulation analysis in Fig. 5, the Anderson estimator showed mixed performance, especially at low sample sizes. However at higher sample sizes, the Anderson estimator was able to improve over collecting additional neural data using just behavioral data. We note that these results do not perfectly match the simulations in the main text, likely for two reasons. First, the correlation estimate we present in parentheses is still estimated from the data and thus is measured with error. We therefore cannot perfect align the simulations with the corresponding true correlation. Second, while we can compute a correlation for arbitrary bivariate data, the actual data may deviate from the bivariate normal used in the simulations. For instance, the distributions in the HCP dataset may be highly skewed. Therefore, we would not expect the simulations to perfectly match these results. We hope that future work will sort out how to improve estimation at reasonable sample sizes.

### A.4 Unbiased Anderson 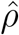

As noted in the main text, the Anderson 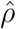 estimator utilizes estimates of the variance of the behavioral variable with the *N*_*xy*_ subjects and the full *N*_*x*_ dataset. The maximum likelihood estimate, normalized by *N* is known to be biased and the unbiased estimate is normalized by *N* − 1. While all previous treatments of Anderson’s *ρ* have used the biased MLE, we analyze the performance of the unbiased estimate in the simulation setup used for Fig. 5. We find that the unbiased estimate performs favorably, with decreased variance in the low *N*, low population correlation regime at the expense of slightly higher variance in the high *N*, high population correlation regime. Given that most cognitive neuroscience studies are likely to be in the former, we recommend using the unbiased variance estimates and indeed that is what we do in all other simulations in the present work.

### A.5 Power for group-level parameters as a function of model fit

From Wilson and Niv (2015), we know the distribution of estimates in the first level regression, i.e.

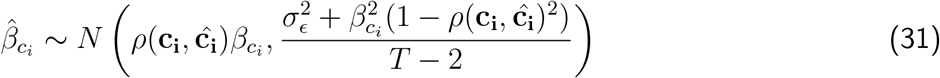

Given the population distributions above and using standard formulas for linear combinations of Gaussian random variables, we can get

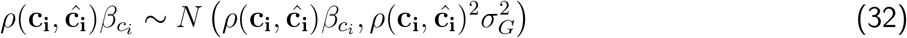

We can now write the distribution of first level estimates as

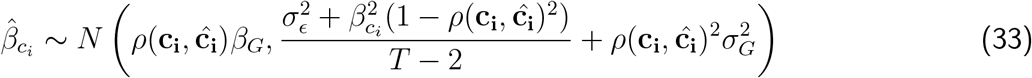

The parameter we are interested in is *β*_*G*_, the average effect in the population. While there are more efficient but computationally expensive estimators (Beckmann et al., 2003; Woolrich et al., 2004), the most common way to estimate *β*_*G*_ is the OLS method, i.e.

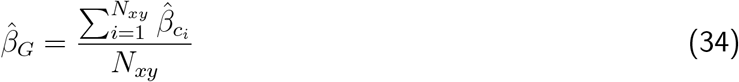

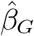 is then just the mean of independent Gaussian random variables and will therefore have a Gaussian distribution with mean *ρ*(**c**_**i**_, **ĉ**_**i**_)*β*_*G*_ and variance

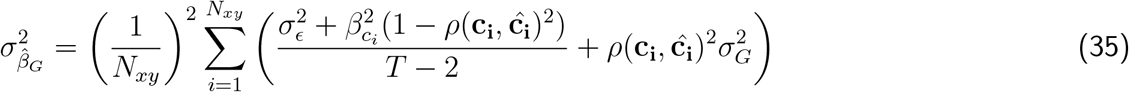

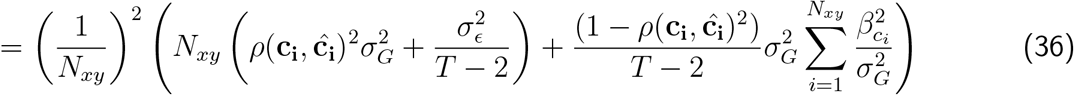

The first term is a constant. In the second term, because *β*_*c*_ is a Gaussian random variable, the normalized squared sum has a non-central chi-squared distribution,

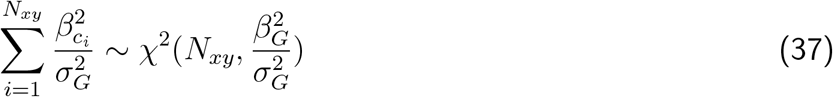

and a non-central chi-squared multiplied by a constant will have a generalized chi-squared distribution. Plugging in the mean of that distribution, the expectation of the variance is now:

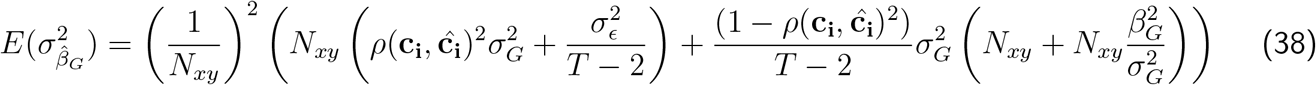

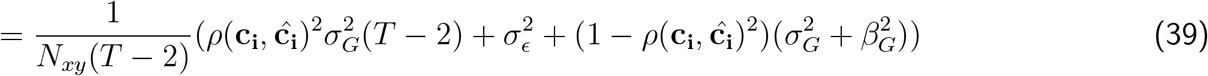

Combining this with the expected estimate, we can get an *t*-statistic at the expected variance

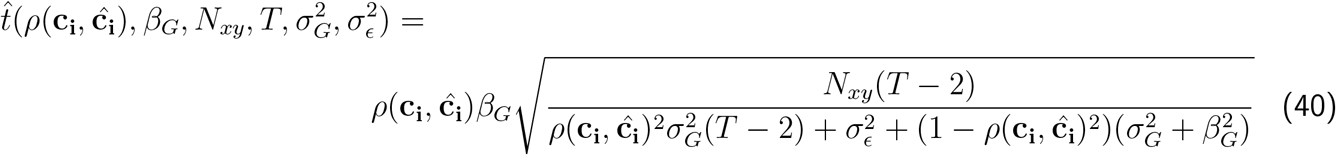

Rather than parameterizing this in terms of the raw effect size, *β*_*G*_, it can be useful for understanding to standardize the variables by the total standard deviation, 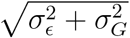, similar to the derivation in Wilson and Niv (2015). For clarity, we define two new variables. Following Wilson and Niv (2015), we call the standardized effect size, 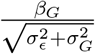, the contrast-to-noise ratio (CNR). In multilevel modeling, an important statistic is the intra-class correlation coefficient or ICC which is the proportion of the total variance that is explained by the subject level variance, i.e. 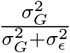 (Chen et al., 2018; Shrout & Fleiss, 1979). If we multiply the right hand side of the above equation by 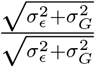, we can now write the equation in terms of these two variables, i.e.,

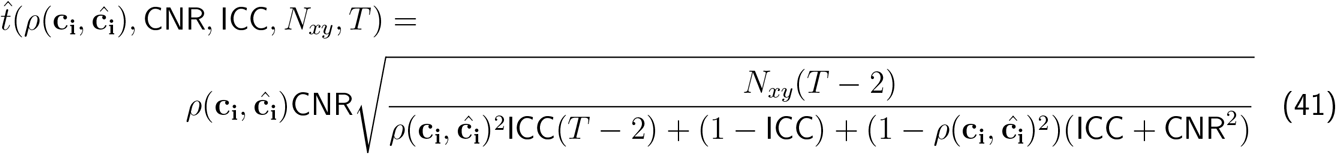

## Footnotes

1 Indeed, it may seem a bit strange to distinguish between these two types of analyses. While conceptually different, researchers frequently use them interchangeably. This is likely due to the fact that standard *t*-tests for regression slopes and correlation coefficients are mathematically identical. However, this is no longer true when we use alternative estimators like the ones described below and focus on precision rather than testing.

2 All figures and simulations in this paper are created using custom code in `python 3.7.7` using the packages `pandas` (McKinney, 2010), `numpy` (van der Walt et al., 2011), `matplotlib` (Hunter, 2007) and `seaborn` (Waskom et al., 2020)

3 In the following we will focus on the case where we are interested in a single latent cognitive variable.

## Notes

### Competing Interest Statement

The authors have declared no competing interest.

### Summary of Updates

Various revisions to text; Several figures revised to have more interpretable axes; New simulations added to appendix.

